# Rational design of hepatitis C virus E2 core nanoparticle vaccines

**DOI:** 10.1101/717538

**Authors:** Linling He, Netanel Tzarum, Xiaohe Lin, Benjamin Shapero, Cindy Sou, Colin J. Mann, Armando Stano, Lei Zhang, Kenna Nagy, Erick Giang, Mansun Law, Ian A. Wilson, Jiang Zhu

## Abstract

Hepatitis C virus (HCV) envelope glycoproteins E1 and E2 are critical for cell entry with E2 being the major target of neutralizing antibodies (NAbs). Here, we present a comprehensive strategy for B cell-based HCV vaccine development through E2 optimization and nanoparticle display. We redesigned variable region 2 in a truncated form (tVR2) on E2 cores derived from genotypes 1a and 6a, resulting in improved stability and antigenicity. Crystal structures of three optimized E2 cores with human cross-genotype NAbs (AR3s) revealed how the modified tVR2 stabilizes E2 without altering key neutralizing epitopes. We then displayed these E2 cores on 24- and 60-meric nanoparticles and achieved high yield, high purity, and enhanced antigenicity. In mice, these nanoparticles elicited more effective NAb responses than soluble E2 cores. Next-generation sequencing (NGS) defined distinct B-cell patterns associated with nanoparticle-induced antibody responses, which cross-neutralized HCV by targeting the conserved neutralizing epitopes on E2.

**One Sentence Summary:** An HCV vaccine strategy is presented that displays redesigned E2 cores on nanoparticles as vaccine candidates for eliciting a broadly neutralizing B-cell response.

---

Hepatitis C virus (HCV) infects 1-2% of the world population and poses a major health burden that leads to ∼500,000 deaths annually and an estimated 1.5-2 million new infections each year (*1, 2*). The opioid epidemic, causing over 70,000 overdose-related deaths in 2017 alone (*3*), is directly contributing to the rapid rise of HCV infection in North America (*4*). Most HCV patients (75– 85%) will develop a chronic infection resulting in hepatocellular carcinoma, cirrhosis, and other severe liver diseases (*1*). Although direct-acting antiviral (DAA) therapies have increased the HCV cure rate (*5, 6*), challenges remain because diagnosis often occurs at a late stage after liver damage (*7*). DAA treatment cannot prevent HCV reinfection nor reduce the risk of liver cancer in advanced liver disease (*8–10*) and resistance may emerge. Indeed, increased HCV-associated mortality and new infections in injection drug users (IDU) (*4, 11, 12*) highlights the urgent need to develop an effective prophylactic vaccine to combat HCV.

A major challenge in HCV vaccine development is how to elicit a broadly protective immune response to overcome the high genetic diversity of six major HCV genotypes and more than 86 subtypes (*13*). Moreover, rapid mutation leads to viral quasispecies in infected individuals that result in immune escape (*14*). Notwithstanding, spontaneous viral clearance in 20-30% of acutely infected patients suggests that chronic HCV infection is preventable if an effective immune response can be induced by vaccination. Glycoproteins E1 and E2 form a heterodimer on the HCV envelope that mediates viral entry into host hepatocytes (*15*). E2 interacts with host cellular receptors CD81 and SR-B1 (*16*) and is a major target for neutralizing antibodies (NAb) (*17*). Crystal structures of an E2 core (E2c) from isolate H77 (genotype 1a) with a broadly neutralizing antibody (bNAb), AR3C, and a truncated E2 from isolate J6 (genotype 2a) bound to a non-NAb, 2A12, provided the first insight into immune recognition of HCV envelope glycoproteins and paved the way for structure-based design of antiviral drugs and vaccines (*18, 19*). Diverse vaccine strategies such as viral vectors, DNA vaccines, virus-like particles (VLP), and recombinant E2 and E1E2 proteins have been explored (*20*), but no licensed vaccine is available to prevent HCV infection. Although recombinant E1, E2 and E1E2 glycoproteins have elicited NAbs in animals and humans (*21*), neutralization breadth was limited and directed mainly to the immunodominant variable loops. Therefore, HCV vaccine efforts should be focused on design and optimization of envelope glycoprotein-based antigens capable of eliciting a bNAb response.

Over the last decade, several rational vaccine design strategies for human immunodeficiency virus type-1 (HIV-1) have included epitope-focused (*22, 23*) and native Env trimer-based approaches (*24, 25*), that aim to direct the immune response to bNAb epitopes either by grafting the epitope onto heterologous scaffolds, removing or suppressing immunodominant regions, or stabilizing Env structures. Another major advance was the development of self-assembling nanoparticles (NP) to present epitope-scaffolds and stabilized Env trimers as multivalent VLP vaccines (*26–31*). These general design elements can, in principle, be applied to a wide range of vaccine targets including HCV. Indeed, epitope-scaffolds have been designed for conserved E1 and E2 NAb epitopes (*32–34*), but with no reported *in vivo* data or little improvement in neutralization breadth. Recent crystal structures of partial E2 ectodomain (E2_ECTO_) – without hypervariable region 1 (HVR1) and stalk – in complex with NAbs HEPC3 and HEPC74 indicate that variable loops may occlude antibody access to conserved neutralizing epitopes on E2 or E1E2 interface (*35*). Here, we designed a truncated variable region 2 (tVR2) on E2 cores from genotypes 1a and 6a and displayed them on self-assembling nanoparticles to assess their vaccine potential.

## Structure-based optimization of HCV envelope glycoprotein E2 core

The HCV E2_ECTO_ is stabilized by nine conserved disulfide bonds, contains three variable regions including HVR1 (a.a. 384-410), VR2 (a.a. 460-485), and VR3 (a.a. 572-597), and is covered with ∼11 *N*-linked glycans (*36*) (Fig. 1A). HVR1 modulates SR-BI interaction (*37*) and all variable regions facilitate host immune evasion by generating escape mutations and shielding neutralizing epitopes (*38, 39*). Empirical engineering (fig. S1A) enabled E2 structure determination (*18, 19*) by shortening the N-/C-termini and VR2, removing glycans at N448 and N576 (*18*), and further V3 truncation (*40, 41*). The E2 core contains an immunoglobulin (Ig)-like β-sandwich domain with front and back layers (*18*) (fig. S1B). The CD81 receptor binding site is a hydrophobic patch formed by the front layer and CD81 binding loop, and overlaps the E2 neutralizing face (*42*) (fig. S1B, middle). However, current E2c constructs exhibit high flexibility involving the front layer C-terminus, shortened VR2, and β-sandwich N-terminus.

**Fig. 1.**
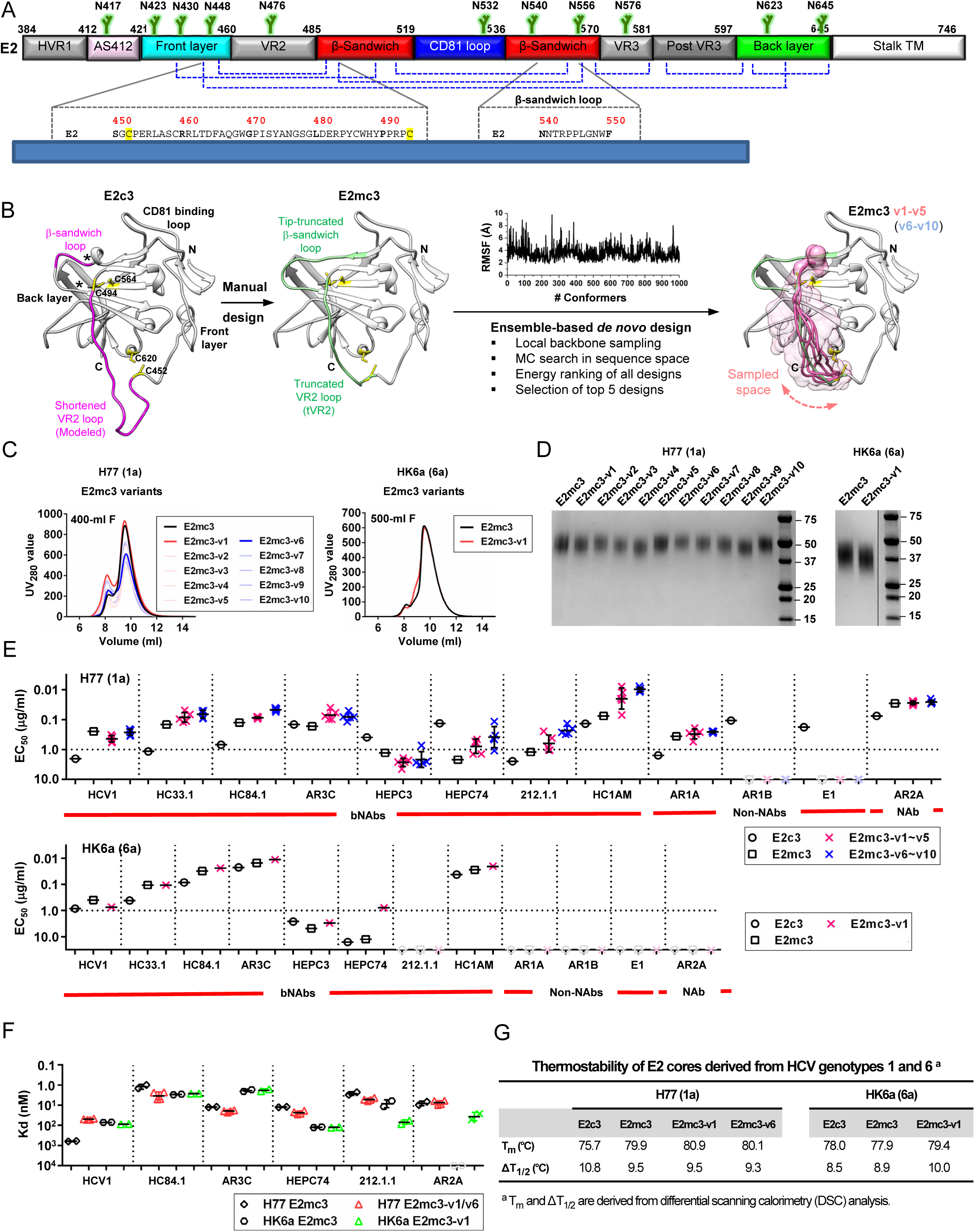
Rational design of HCV E2 cores. (**A**) Schematic representation of HCV E2 (amino acids 384-746) colored by structural components with variable regions (VRs) in gray, antigenic site 412 (AS412) in pink, front layer in cyan, β-sandwich in red, CD81 binding loop in blue, back layer in green, stalk trans-membrane (TM) region in white, and *N*-linked glycans and conserved disulfide bonds are indicated by green branches and blue dashed lines respectively. Sequence alignment of the design regions between E2 and E2mc3 is shown below. **(B)** Structure-based design of E2 mini-cores. Left: Structure of H77 E2c3 (modeled upon H77 E2c in PDB ID: 4MWF) with shortened VR2 loop modeled by LOOPY. The redesigned β-sandwich loop and the shortened VR2 loop are colored in magenta. Disulfide bonds, C494-C564 and C452-C620, which anchor the VR2 loop to the back layer, are shown in yellow sticks. Front layer, CD81 binding loop, and back layer are also labeled. Middle 1: Structure of H77 E2mc3 with tip-truncated β-sandwich loop and further truncated VR2 loop (tVR2) are colored in green. Middle 2: root-mean-square fluctuation (RMSF) plot for redesigned tVR2 ensemble is shown with the major steps involved in the ensemble-based *de novo* protein design below. Right: Structure of H77 E2mc3 with five top-ranking tVR2 design variants (E2mc3 v1-v5) colored in pink and highlighted in a transparent molecular surface. **(C)** SEC profiles of E2mc3 and variants. Left: H77 E2mc3 (in black), v1-v5 (v1 in red and v2-v5 in light red), and v6-v10 (v6 in blue and v7-v10 in light blue). Right: HK6a E2mc3 (in black) and v1 (in red). (**D**) SDS-PAGE of E2mc3 and variants (Left: H77; Right: HK6a). (**E**) EC_50_ (μg/ml) values of H77 (upper panel) and HK6a (lower panel) E2 cores binding to 12 HCV antibodies, including eight bNAbs (HCV1, HC33.1, HC84.1, AR3C, HEPC3, HEPC74, 212.1.1, and HC1AM), one NAb (AR2A), and three non-NAbs (AR1A, AR1B, and E1). E2 cores tested here include E2c3, E2mc3, and E2mc3 variants (10 for H77 and 1 for HK6a). **(F)** Binding affinities (Kds, in nM) of H77 and HK6a E2mc3 variants for six selected HCV antibodies. **(G)** Thermal stability of H77 and HK6a E2c3 and E2mc3 variants measured by DSC. Two thermal parameters, *T_m_* and ΔT_1/2_, are listed for four H77 E2 cores and three HK6a E2 cores.

Here, we redesigned the VR2 disordered region that is anchored to the back layer and β-sandwich by two disulfide bonds, C452-C620 and C494-C564 (Fig. 1, A and B). Although this region consists of 43 residues in wild-type E2 and 21 residues in E2c/E2c3, the Cα distance between C452 and C494 is 26.3Å for H77 E2c (fig. S1B), which could be jointed using a minimum of 7 residues. We first manually truncated the VR2 loop region to 13 residues (tVR2) (Fig. 1, A and B; fig. S1A) and deleted the tip (aa 543-546) of the β-sandwich loop (a.a. 540-552) to focus the immune response to bNAb epitopes, as non-Nabs, such as AR1B, E1 and HEPC46, bind to this region (*40, 43, 44*) (figs. S1C-E). The new E2 core is termed E2 mini-core 3 (E2mc3). Next, we redesigned tVR2 in H77 E2mc3 for two loop lengths, 13 aa (as in E2mc3) and 12 aa, using ensemble-based *de novo* protein design (*45*) to identify optimal tVR2 sequences that stabilize E2mc3 (Fig. 1B). For each tVR2 length, an ensemble of loop conformations (1,000) was generated to connect C452 and C494 (fig. S1F) with Cα root-mean-square fluctuation (RMSF) ranging from 1.9 to 9.8 Å (average 3.6 Å) and 1.6 to 7.8 Å (average 3.2 Å) for 13 aa and 12 aa loops, respectively (fig. S1G). After extensive Monte Carlo sampling, the five top-ranking sequences for each loop length, E2mc3-v1-v5 and E2mc3-v6-v10 (Fig. 1B, right; fig. S1H), were selected for further characterization. As HK6a E2c3 and H77 E2c share high structural similarity and disordered regions when bound to AR3 bNAbs (*40*), we designed HK6a E2mc3 and E2mc3-v1 constructs (fig. S1I) without further modification. In total, eleven H77 E2 cores and two HK6a E2 cores were advanced to experimental evaluation.

## Biochemical, biophysical, and antigenic assessment of HCV E2mc3 designs

Previously, we extensively characterized our rationally designed HIV-1 trimers and nanoparticles to facilitate immunogen selection for *in vivo* testing (*26, 27, 45*). Here, 13 E2mc3 constructs from H77 and HK6a and two parental E2c3 constructs (*40, 41*) were transiently expressed in HEK293 F cells and purified using immunoaffinity (*18*) followed by size exclusion chromatography (SEC). Overall, the purified E2mc3 variants showed greater yield than their respective E2c3 constructs ranging from 5.0 to 11.5 mg from 1L HEK293 F transfection. AR3A-purified E2mc3 was mostly in monomeric form with a small aggregate peak (Fig. 1C and fig. S2A) and SEC-purified proteins ran as a single band (∼50 kDa) on SDS-PAGE (Fig. 1D and fig. S2B). We then tested H77 and HK6a E2mc3 variants by ELISA on a panel of HCV antibodies that included (b)NAbs targeting antigenic site 412 (AS412), AS434, antigenic region 3 (AR3), AR2 (*42*), and non-NAbs targeting AR1 (fig. S2C). H77 E2mc3 showed greater binding than E2c3 for most bNAbs (excluding HEPC3/74 (*46*)) and NAb AR2A, with further improvements for some E2mc3 variants (Fig. 1E, upper panel; fig. S2, D and E). As expected, truncation of the β-sandwich loop reduced binding to non-NAbs AR1B and E1 with negligible effect on AR1A, which recognizes AR1 but not the β-sandwich loop (*43*). Similar patterns were observed for HK6a E2mc3 and E2mc3-v1 except for NAb 212.1.1 (*47*) (Fig. 1E, lower panel; fig. S2, F and G) with no detectable binding by genotype-specific NAbs AR2A and AR1A/B (*43*). By biolayer interferometry (BLI), HK6a E2mc3 variants exhibited similar antigenic profiles (Fig. 1F; figs. S2, H to J). Differential scanning calorimetry (DSC) showed a 4.2°C increase in *T_m_* for E2mc3 (H77) relative to E2c3 that was further increased by 1°C and 0.2°C for E2mc3-v1 and v6, respectively (Fig. 1G and fig. S2K) (*48*).

## Structural characterization of minimized cores derived from H77 and HK6a

Crystallization of H77 and HK6a E2 cores with antigen-binding fragments (Fabs) derived from AR3A/B/C/D bNAbs led to structures of H77 E2mc3-v1 and E2mc3-v6 with AR3C and HK6a E2mc3-v1 with AR3B at 1.90 Å, 2.85 Å, and 2.06 Å, respectively (Fig. 2A and table S2). The overall fold of E2mc3 variants is highly similar to H77 E2c and HK6a E2c3 (PDB: 4MWF and 6BKB) (Fig. 2A), but with differences in the back layer C-terminus (a.a. 629-640) (*40*) and a front layer loop (a.a. 430-438) (*18*) (fig. S3A) that interacts with heavy-chain complementarity-determining region 3 (HCDR3) of bNAbs AR3A-D. However, similar hydrophilic contacts are maintained with HCDR3 (fig. S3A) further supporting the conformational plasticity of the E2 front layer (*41*). The shortened β-sandwich loop can be fully modeled in the H77 E2mc3-v1 complex with bNAb AR3C (fig. S3B), confirming its truncation results in loss of key interactions with non-NAb E1 (Fig. 1E and Fig. 2B). The redesigned tVR2 can be fully modeled in AR3s-bound H77 and HK6a E2mc3-v1 structures, but is only partially visible in AR3C-bound H77 E2mc3-v6 (Fig. 2C and fig. S4A). The tVR2 redesign does not introduce any conformational changes to the E2 neutralizing face (Fig. 2D), which is anchored to the back layer and β-sandwich by C452-C620 and C494-C564 and interacts with the truncated VR3 and β-sandwich (fig. S4B). While H77 and HK6a E2mc3-v1 constructs share the same tVR2 sequence, a significant difference in conformation (Fig. 2C and fig. S4C) likely results from differences in sequence and structure of the adjacent VR3 and β-sandwich loop (*40*) in genotypes 1 and 6 (figs. S3B and S4C). The tVR2 redesign has also minimal effect when compared to the recent genotype-1 E2_ECTO_ structures in complex with bNAbs HEPC3/74 (*35*) (figs. S1A and S4D).

**Fig. 2.**
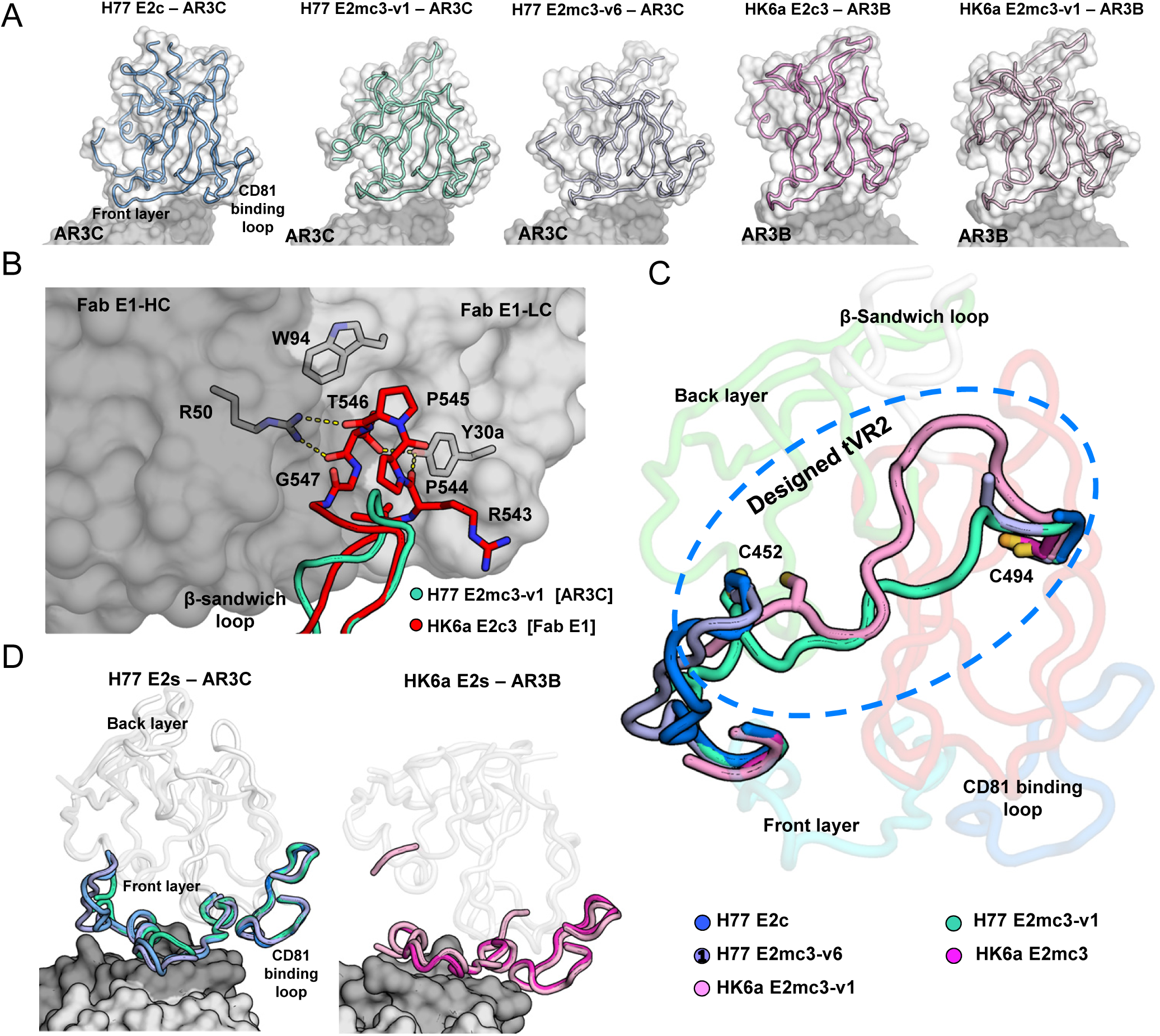
Structures of rationally designed HCV E2 cores. **(A)** Crystal structures of H77/HK6a E2mc3 indicate an overall similar fold to H77 E2c and HK6a E2c3 (PDB: 4MWF and 6BKB). **(B)** Superposition of the β-sandwich loop from the H77 E2mc3-v1 structure on the HK6a E2c3-Fab E1 complex confirming that loss of binding of E2mc3s to Fab E1 results from truncation of the β-sandwich loop. **(C)** Superposition of E2 of HK6a E2c3 (PDB 6BKB), H77 E2mc3-v1, H77 E2mc3-v6, and HK6a E2mc3-v1 on the structure of H77 E2c (PDB 4MWF) illustrating the conformation of the redesigned tVR2 (a.a. 452-494). The redesigned tVR2 regions of H77 E2mc3-v1 and HK6a E2mc3-v1 structures are fully modeled but only partly in the H77 E2mc3-v6 structure. **(D)** Superposition of the H77/HK6a E2mc3 structures to H77 E2c and HK6a E2c3 indicating similar conformation of the neutralization face with only local conformational changes for the redesigned VR2 E2s.

## Design and characterization of nanoparticles presenting optimized E2 cores

Improved immunogenicity in mice was recently reported for a ferritin nanoparticle carrying soluble E2 (sE2, a.a. 384-661) that contains three full-length immunodominant variable loops (*49*). Here, we displayed our E2mc3 variants, which only present the conserved bNAb epitopes, on self-assembling nanoparticles as multivalent HCV vaccine candidates (Fig. 3A). We tested three nanoparticle platforms: 24-meric ferritin (FR, as control) and 60-meric E2p and I3-01, ranging in size from 24.5-37.5 nm (Fig. 3B). We genetically fused the C-terminus of E2mc3-v1 to the N-terminus of the nanoparticle subunit via a 10-residue linker, (G_4_S)_2_, termed 10GS. Six constructs were transiently expressed in ExpiCHO or 293 F cells and purified on an AR3A column followed by SEC (Fig. 3C and fig. S5A). For H77, the SEC profiles demonstrated substantial yield and purity for all E2 core nanoparticles with different patterns for 24- vs. 60-mers (Fig. 3C). For HK6a, reduced nanoparticle yield and purity was accompanied by increased low-molecular-weight species (fig. S5A), suggesting H77 tVR2 may be less compatible with HK6a and hinder particle assembly. However, effective particle assembly was observed in blue-native PAGE (BN-PAGE) and negative-stain electron microscopy (EM) (Fig. 3, D and E; fig. S5, B and C). Enhanced bNAb binding was seen with H77 nanoparticles (up to 100-fold EC_50_ change) and no binding to non-NAbs targeting the β-sandwich loop (Fig. 3F and fig. S5, D and E). HK6a E2mc3-v1 nanoparticles exhibited similar but genotype-specific profiles (Fig. 3F and fig. S5, F and G). In BLI, correlation between peak signal and antigen valency was observed irrespective of genotype with 60-mers > 24-mer > E2 core monomer (Fig. 3G; fig. S5, H and I), consistent with our HIV-1 gp140 nanoparticles (*26, 27*).

**Fig. 3.**
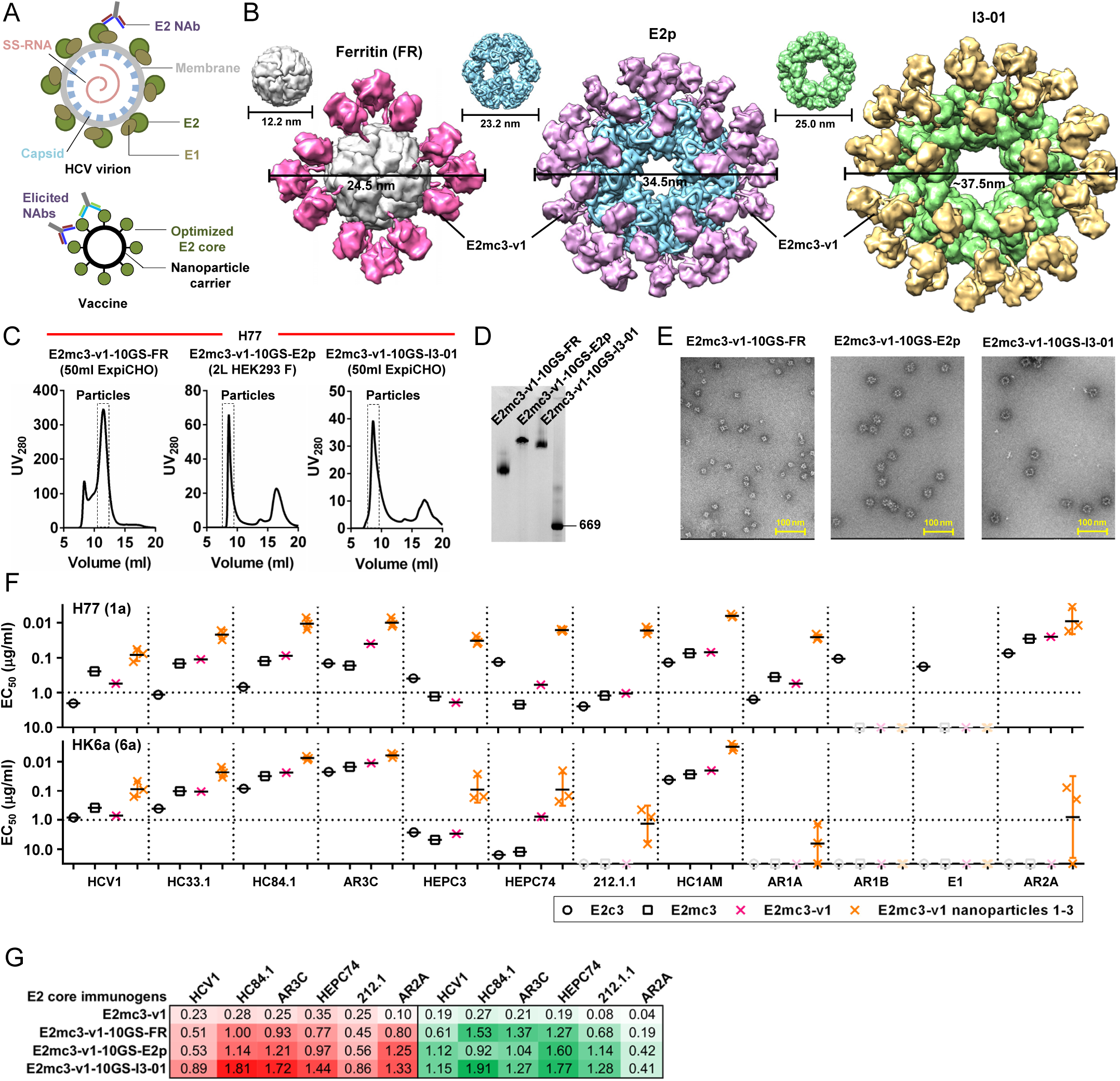
Rational design of self-assembling E2 core nanoparticles. **(A)** Schematic representation of HCV virion (top) and E2 core-based nanoparticle vaccine (bottom). For the HCV virion, single-stranded (SS)-RNA, capsid, membrane, envelope glycoproteins E1 and E2 are labeled, while for the vaccine, optimized E2 core and nanoparticle carrier are labeled. **(B)** Colored surface models of nanoparticle carriers (top) and E2 core-based nanoparticle vaccines (bottom). Three nanoparticle carriers shown here are 24-meric ferritin (FR) and 60-meric E2p and I3-01. Nanoparticle size is indicated by diameter (in nanometers). **(C)** SEC profiles of H77 E2mc3-v1 nanoparticles obtained from a Superose 6 10/300 GL column. The particle fraction is indicated by a dotted-line box. While both FR and I3-01 nanoparticles were produced in ExpiCHO cells, E2p nanoparticles were expressed in HEK293 F cells. **(D)** BN-PAGE of SEC-purified H77 E2mc3-v1 nanoparticles. **(E)** Negative stain EM images of SEC-purified H77 E2mc3-v1 nanoparticles. (**F**) EC_50_ (μg/ml) values of H77 (upper panel) and HK6a (lower panel) E2mc3-v1 nanoparticles binding to 12 HCV antibodies listed in Fig. 1C. **(G)** Antigenic profiles of H77 (left, in red) and HK6a (right, in green) E2mc3-v1 and three nanoparticles against six HCV antibodies. Sensorgrams were obtained from an Octet RED96 using an antigen titration series of six concentrations (3.57-0.11 μM by twofold dilution for E2mc3-v1 and 52.08-1.63 nM by twofold dilution for nanoparticles) and quantitation biosensors, as shown in fig. S5, H and I. The peak siginals (nm) at the highest concentration are listed in the matrix. Higher color intensity indicates greater binding signal measured by Octet.

## E2 core nanoparticles elicit stronger immune responses than E2 cores in mice

We assessed H77 and HK6a E2 core nanoparticles in wild-type BALB/c mice in studies #1 and #2, respectively (*50*), using a short regimen (Fig. 4A). In study #1, three H77-based vaccines showed a correlation between E2-specific EC_50_ titer and antigen valency at week 2 with significant *P*-values (Fig. 4B, upper panel; fig. S6, A and B). While E2-specific antibody titers continued to rise, the difference between the three vaccine groups diminished at week 11. In study #2, we compared HK6a E2mc3-v1, its E2p nanoparticle, and HK6a/H77 E2mc3-v1 E2p nanoparticle mix (Fig. 4B, lower panel; fig. S6, C and D). The HK6a E2mc3-v1 E2p group retained its advantage in antibody titer until week 8, whereas its H77 counterpart did until week 11. The E2p mix elicited significantly higher titers to H77 than to HK6a throughout the immunization. Overall, E2 core nanoparticles induced greater antibody titers than E2 cores, although E2 only accounts for 42% (E2p) to 51% (FR) of the protein mass. We then evaluated serum neutralization using HCV pseudoparticles (HCVpp) (*43*). In study #1, autologous neutralization increased steadily over time with distinct temporal patterns (Fig. 4C, upper panel). At week 2, the FR group showed the highest H77 neutralization, whereas the E2p group was unexpectedly the lowest (Fig. 4C, upper panel). From week 5, the FR grouped show lower neutralizing activity with a significant *P*-value at week 11, whereas the E2p group became the best performer with statistical significance at weeks 8 and 11. Week 11 sera also neutralized heterologous isolates HCV-1 (1a), J6 (2a), and SA13 (5a), with significant *P*-values for HCV-1 and J6 (Fig. 4C, lower panel). A similar trend was observed for study #2 (Fig. 4D) (*51*), and the E2p mix group was equivalent to the HK6a-only E2p group. Five HCV bNAbs and HIV-1 bNAb VRC01 (*52, 53*) validated the HCVpp assays (Fig. 4E).

**Fig. 4.**
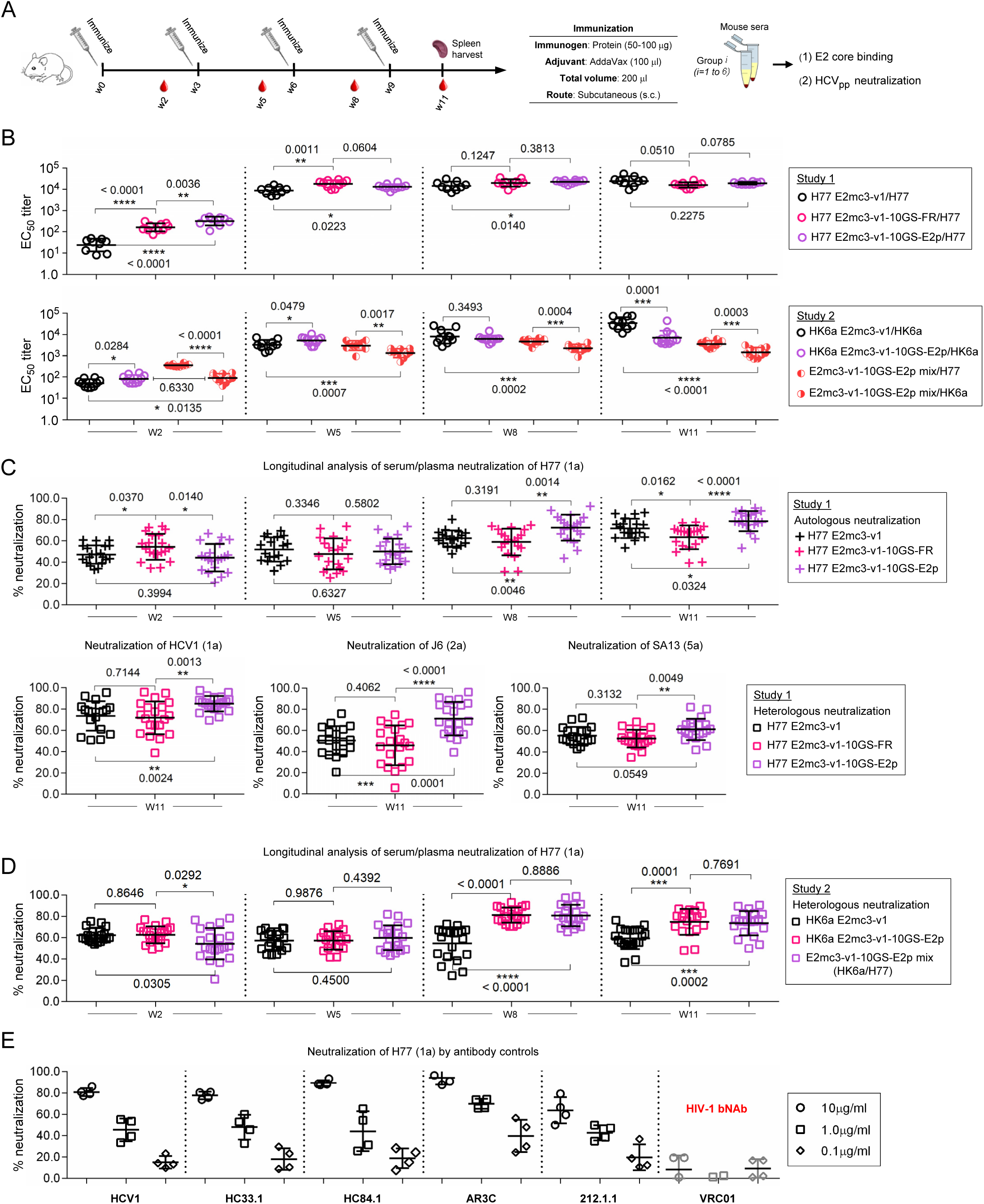
Immunogenicity of newly designed E2 cores and nanoparticles in mice. **(A)** Schematic representation of the mouse immunization protocol. In study #1, mice were immunized with H77 E2mc3-v1 (group 1), H77 E2mc3-v1-10GS-FR (group 2), and H77 E2mc3-v1-10GS-E2p (group 3). In study #2, mice were immunized with HK6a E2mc3-v1 (group 1), HK6a E2mc3-v1-10GS-E2p (group 2), and HK6a/H77 E2mc3-v1-10GS-E2p mix (group 3). **(B)** Longitudinal analysis of E2-specific antibody titers in immunized mouse sera at weeks 2, 5, 8 and 11. Top panel: EC_50_ titers (fold of dilution) calculated from ELISA binding of mouse sera in study #1 to the coating antigen, H77 E2mc3-v1. Bottom panel: EC_50_ titers calculated from ELISA binding of mouse sera in study #2 to the coating antigens HK6a E2mc3-v1 (groups 1-3) and H77 E2mc3-v1 (group 3). The *P*-values were determined by an unpaired *t* test in GraphPad Prism 6 and are labeled on the plots, with (*) indicating the level of statistical significance. Detailed serum ELISA data is shown in figs. S6, A to D. **(C)** Mouse serum neutralization in study #1. Top panel: Percent (%) neutralization of mouse sera against autologous H77 at weeks 2, 5, 8 and 11. Bottom panel: Percent (%) neutralization of mouse sera against heterologous HCV-1, J6, and SA13 at the last time point, week 11. **(D)** Mouse serum neutralization in study #2. Percent (%) neutralization of mouse sera against heterologous H77 at weeks 2, 5, 8 and 11. **(E)** Validation of the HCV pseudotyped particle (HCVpp) neutralization assay using five HCV bNAbs and an HIV-1 bNAb (negative control) against H77. Percent (%) neutralization of all antibodies was determined at three concentrations, 10.0μg/ml, 1.0μg/ml, and 0.1μg/ml.

## Distinctive patterns of B cell responses induced by E2 core and E2 core nanoparticle

We combined antigen-specific B cell sorting and antibody NGS to obtain a quantitative readout of vaccine-induced B cell responses and determine B cell patterns associated with different vaccine platforms (Fig. 5A). We used an H77 E2mc3-v1 probe with an C-terminal Avi-tag (fig. S7A) to sort E2-specific splenic B cells from mice in the H77 E2mc3-v1 and E2p nanoparticle groups by flow cytometry. A greater frequency/number of E2-specific B cells was observed for the E2p group with significant *P*-values (Fig. 5B and fig. S7B). Sorted B cells from 10 mice, five per group, were subjected to NGS (*40*) and repertoire analysis (*26, 29*) (fig. S7C). The E2p group used significantly more (5 to 8) heavy-chain variable (V_H_) genes than the E2 core group (∼1) in each animal (Fig. 5C). Antibodies elicited by E2p contained more V_H_ mutations with a significant *P*-value of 0.0268 (Fig. 5D). Distinct patterns of HCDR3 length were observed for the two groups with the E2 core group showing two dominant HCDR3 lengths, while the E2p group produced a much broader distribution (Fig. 5E). The E2p group yielded a greater average RMSF – the range in which the loop length varies – than the E2 core group (4.5aa vs. 1.1aa) with a *P*-value of <0.0001.

**Fig. 5.**
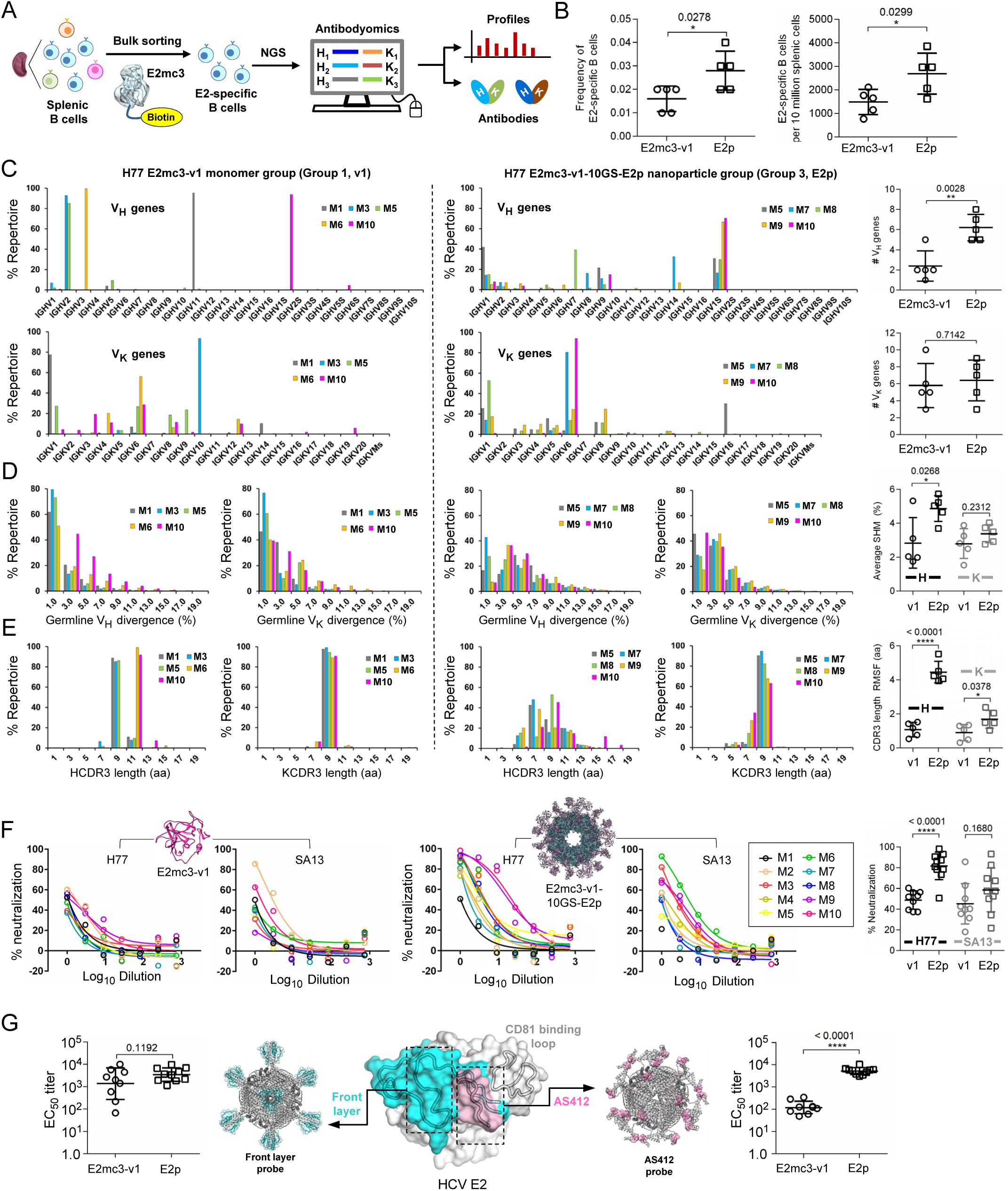
Patterns associated with HCV E2-specific B cell response in mouse immunization. **(A)** Schematic representation of the strategy used to analyze HCV E2-specific B cell response that combines antigen-specific bulk sorting of splenic B cells with next-generation sequencing (NGS) and antibodyomics analysis. **(B)** Statistical analysis of B cell sorting data obtained for group 1 (H77 E2mc3-v1 monomer) and group 3 (H77 E2mc3-v1-10GS-E2p nanoparticle) in study #1. Left: Frequency of E2-specific B cells. Right: Number of E2-specific B cells per million splenic cells. Five mice from group 1 (M1, M3, M5, M6 and M10) and five mice from group 3 (M5, M7, M8, M9, and M10) were randomly selected and analyzed. **(C)** Distribution of germline gene usage plotted for group 1 and group 3. Top panel: Germline V_H_ genes. Bottom panel: Germline V_κ_ genes. Statistical analysis of number of activated V_H_/V_κ_ genes (≥1% of the total population) is shown on the far right. **(D)** Distribution of germline divergence or degree of somatic hypermutation (SHM) plotted for groups 1 and 3. For each group, percent (%) mutation is calculated at the nucleotide (nt) level for V_H_ (left) and V_κ_ (right). Statistical analysis of germline divergence is shown on the far right. **(E)** Distribution of CDR3 loop length plotted for groups 1 and 3. For each group, CDR3 length calculated at the amino acid (a.a.) level is shown for heavy (left) and light chains (right). Statistical analysis of root-mean-square fluctuation (RMSF) of CDR3 loop length, which is used as an indicator of how much the CDR3 loop length varies within the E2-specific antibodies from each animal. **(F)** Neutralization curves using purified IgG for groups 1 (left) and 3 (right) in study #1. Autologous H77 (1a) and heterologous SA13 (5a) were tested in HCVpp assays with a starting IgG concentration of 100μg/ml followed by a series of three-fold dilutions. Structural models of the immunogens are placed next to their neutralization curves. **(G)** Epitope mapping of polyclonal antibody sera from groups 1 and 3 in study #1. Surface model of E2 ectodomain (E2_ECTO_) is shown in the middle with the front layer (FL) and AS412 colored in cyan and pink, respectively. Statistical analysis of EC_50_ titers (fold of dilution) of groups 1 and 3 against the FL probe (left) and the AS412 probe (right). Structural models of the designed nanoparticle probes are placed next to their plots. Epitopes on the nanoparticles are colored according to the E2_ECTO_ model.

## Polyclonal NAbs induced by E2 core and E2 nanoparticle target different epitopes

Mouse serum contains nonspecific antiviral activity, which may interfere with HCVpp assays (*54*). We purified mouse IgG from study #1 at week 11 for neutralization of H77 and SA13 HCVpps with starting IgG concentration at 100 μg/ml followed by a series of three-fold dilutions (Fig. 5F and fig. S8A). For H77, while no mice sera in the E2 core group exhibited >60% neutralization at the first concentration, mice #9 and #10 in the E2p group showed plateaued curves, suggesting potent NAbs in the IgG, with a similar but less pronounced trend for SA13. Unpaired *t* tests indicated a significant difference between the E2p and E2 core groups for H77 (*P*<0.0001), but not for SA13 (*P*=0.1680). The FR group ranked the lowest in serum neutralization but slightly outperformed the E2 core group in IgG neutralization (fig. S8A). Nonetheless, ferritin may not be an optimal platform for HCV nanoparticle vaccine design. Here, two epitope-specific probes were used to examine vaccine-induced antibody responses to two prominent bNAb epitopes: front layer (FL, a.a. 421-459), integral to the E2 neutralizing face (*42*), and AS412 (Fig. 1A and Fig. 5G, middle). A trimeric scaffold was designed to present FL, which was anchored to each subunit via an engineered disulfide bond (fig. S8B). This trimer FL-scaffold was displayed on FR. In ELISA, the E2p group yielded an average EC_50_ titer of 4281 compared to 3044 for the E2mc3-v1 group (Fig. 5G, left and fig. S8C). However, unpaired *t* test reported a non-significant *P*-value, 0.1192, between the two groups (Fig. 5G, left). Nonetheless, nanoparticle display improved recognition of FL including antigenic site 434 (AS434, a.a. 434-446). We then used a previously designed FR nanoparticle (*32*) (fig. S8B, bottom) to probe AS412-specific response. In ELISA, the E2p group demonstrated a uniform, robust response to the AS412 β-hairpin with average EC_50_ titer of 5584, which is 38-fold greater than the E2mc3-v1 group with a *P*-value of <0.0001 (Fig. 5G, right and fig. S8C, bottom). Thus, particulate display focuses the response on conserved bNAb epitopes. These epitope probes also provide valuable tools for assessment of HCV vaccine candidates.

## DISCUSSION AND FUTURE DIRECTIONS

The recent DAA therapy for chronic HCV infection has raised questions about the necessity of developing an HCV vaccine. However, issues in DAA treatment have begun to surface and a prophylactic vaccine is still required to control HCV transmission (*20, 55*). Although HCV genetic diversity poses a significant challenge, recent advances in E2 structures and HCV bNAbs have paved the way for new B cell-based vaccine strategies (*20, 35, 40, 42, 56, 57*). Here, we redesigned the E2 core constructs (*18, 40, 41*) for genotypes 1a and 6a by truncating VR2 and the β-sandwich loop and optimized the truncated VR2 (tVR2) computionally. These new E2 cores were displayed on nanoparticles of various sizes, showing high yield, high purity, and enhanced antigenicity. Mice were immunized with these constructs and longitudinal serum analysis not only confirmed the superior immunogenicity of E2p-based vaccine constructs but also indicated that ferritin might not be as suitable a vaccine carrier as used recently for E2 ectodomain (*49*). NGS profiling of E2-sorted B cells provided much needed insights as to how an effective nanoparticle vaccine can elicit bNAbs with diversified germline gene usage, accelerated antibody maturation, and expanded range of HCDR3 loop length. Serum analysis with novel probes revealed how particulate display impacts epitope recognition by redirecting antibody responses. Unique to this study, statistical analysis was extensively used to validate the *in vivo* data, providing a rigorous foundation for future comparison of different types of HCV vaccine candidates.

Future investigation should be directed toward several possibilities. First, the suboptimal *in vitro* and *in vivo* data observed for HK6a E2mc3-v1, which bears H77 tVR2, suggests that tVR2 may need to be redesigned for each HCV isolate in the vaccine. Second, despite poor yield and purity when displaying HK6a E2mc3-v1, the I3-01 constructs showed greater antigenicity than their E2p counterparts and could be produced in GMP CHO cells, suggesting that I3-01 may still be a valid nanoparticle display platform for HCV vaccine design, consistent with its outstanding immunogenicity in our previous HIV-1 study (*26*). Lastly, different adjuvants can be tested to ensure the optimal immune outcome. Thus, nanoparticles presenting optimized E2 cores of diverse genotypes, in a mixed form (cocktail or mosaic (*58*)) or sequentially, can now be used to generate rapid, broadly protective NAb responses by targeting conserved E2 epitopes.

## Acknowledgements

This work was funded in part by HIV Vaccine Research and Design (HIVRAD) program (P01 AI124337) (to J.Z.), NIH Grants AI129698 and AI140844 (to J.Z.), UfoVax/SFP-2018-0416 and UfoVax/SFP-2018-1013 (to J.Z.), AI123861 (to M.L. and J.Z.), AI079031 (to M.L.), AI123365 and AI106005 (to M.L. and I.A.W.). We thank Zhenyong Keck and Steven Foung at Stanford University for generous sharing of antibody reagents. We thank R. Stanfield, X. Dai, and M. Elsliger for crystallographic and computational support and H. Tien in the Wilson lab for automated crystal screening. X-ray data sets were collected at the APS beamline 23ID-B (GM/CA CAT) and SSRL beamline 12-2. The use of the APS was supported by the U.S. Department of Energy (DOE), Basic Energy Sciences, Office of Science, under contract DE-AC02-06CH11357. The use of the SSRL Structural Molecular Biology Program was supported by DOE Office of Biological and Environmental Research and by the NIH NIGMS (including P41GM103393) and the National Center for Research Resources (P41RR001209).

## Author contributions

Project design by L.H., N.T., M.L., I.A.W. and J.Z; structural design of E2 cores and E2 core nanoparticles by L.H. and J.Z.; plasmid design and processing by L.H. and C.S.; antigen production, purification, and biochemical characterization by L.H., X.L., B.S., and C.J.M.; HCV antibody production by E.L., N.T. and M.L.; E2 complex crystallization, structure determination, and refinement by N.T. and I.A.W.; DSC measurement by N.T. and I.A.W.; negative-stain EM by L.H. and J.Z.; BLI of E2 cores and E2 core nanoparticles by L.H. and X.L.; mouse serum-antigen ELISA by L.H. and X.L.; mouse serum neutralization by L.H., X.L., and K.N.; antigen-specific mouse B cell sorting by L.H. and L.Z.; mouse B cell sequencing by L.H., X.L., and J.Z.; bioinformatics analysis by L.H., X.L. and J.Z.; antibody neutralization by L.H. and X.L.; IgG purification by L.H. and C.S.; epitope mapping by L.H. and X.L. Manuscript written by L.H., N.T., M.L., I.A.W. and J.Z. All authors were asked to comment on the manuscript. The TSRI manuscript number is 29867.

## Data and materials availability

All data and code to understand and assess the conclusions of this research are available in the main text and supplementary materials. The x-ray coordinates and structure factors have been deposited to the Protein Databank with codes: XXXX etc. Additional data related to this paper may be requested from the corresponding authors.

## Materials and Methods

### Structural design of truncated VR2 (tVR2)

Based on the structure of bNAb AR3C-bound H77 E2c (*18*) (PDB ID: 4MWF), the already shortened VR2 loop in E2c, i.e. the segment between C452 and C494, was manually truncated by removing exposed hydrophobic residues and a disulfide bond (C459-C486), resulting in the H77 E2mc3 construct (fig. S1A). The truncated VR2 (tVR2) was modeled by LOOPY (*59*), a torsion-space loop modeling and prediction program. Computational redesign of tVR2 was then performed using an ensemble-based *de novo* protein design method (*45*) with a focus on the N-terminal region of the peptide sequence PERASGHYPRP between C452 and C494. Briefly, an ensemble of 11-aa (G_6_HYPRP) or 10-aa (G_5_HYPRP) loop conformations was generated to connect C452 and C494 using LOOPY (*59*). For each loop conformation, a starting sequence for the multi-glycine (G_n_) region was selected from a pool of 50 random sequences based on the RAPDF potential (*60*) and subjected to 500 steps of Monte Carlo simulated annealing (MCSA) with the temperature linearly decreasing from 300 to 10K. The lowest-energy sequence for each loop was recorded and all MCSA-derived designs were ranked based on energy at the completion of the process. The top 5 designs from the G_6_ and G_5_ ensembles, termed H77 E2mc3-v1-v5 and v6-v10, respectively, were selected for experimental validation. HK6a E2mc3 and E2mc3-v1 constructs were designed by directly adopting the H77 sequence designs without further modification.

### Expression and purification of E2 antigens

All E2 cores (E2c3, E2mc3, and E2mc3 v1-v10) and E2p-based nanoparticles were transiently expressed in HEK293 F cells (Thermo Fisher) for biochemical, biophysical, and antigenic analyses. Briefly, 293 F cells were thawed and incubated with FreeStyle^TM^ 293 Expression Medium (Life Technologies, CA) in a shaker incubator at 37°C, 135 rpm and 8% CO_2_. When the cells reached a density of 2.0×106/ml, expression medium was added to reduce cell density to 1.0×10^6^ ml^-1^ for transfection with polyethyleneimine (PEI) (Polysciences, Inc). Next, 900 μg of plasmid in 25 ml of Opti-MEM transfection medium (Life Technologies, CA) was mixed with 5 ml of PEI-MAX (1.0 mg/ml) in 25 ml of Opti-MEM. After 30-min incubation, the DNA-PEI-MAX complex was added to 1L 293 F cells. Culture supernatants were harvested five days after transfection, clarified by centrifugation at 1800 rpm for 20 min, and filtered using a 0.45 μm filter (Thermo Scientific). E2 proteins were extracted from the supernatants using an AR3A antibody column as previously described (*18, 40*). Bound proteins were eluted three times, each with 5ml of 0.2 M Glycine (pH=2.2) and neutralized with 0.5ml of Tris-Base (pH=9.0). The proteins were further purified by size exclusion chromatography (SEC) on a Superdex 75 Increase 10/300 GL column (GE Healthcare) for E2 cores and on a Superose 6 10/300 GL column (GE Healthcare) for E2p nanoparticles. E2mc3-v1-attached ferritin and I3-01 nanoparticles were produced in ExpiCHO cells (Thermo Fisher). Briefly, ExpiCHO cells were thawed and incubated with ExpiCHO^TM^ Expression Medium (Thermo Fisher) in a shaker incubator at 37 °C, 135 rpm and 8% CO_2_. When the cells reached a density of 10×10^6^ ml^-1^, ExpiCHO^TM^ Expression Medium was added to reduce cell density to 6×10^6^ ml^-1^ for transfection. The ExpiFectamine^TM^ CHO/plasmid DNA complexes were prepared for 100-ml transfection in ExpiCHO cells following the manufacturer’s instructions. For these two nanoparticles, 100 μg of plasmid and 320 μl of ExpiFectamine^TM^ CHO reagent were mixed in 7.7 ml of cold OptiPRO™ medium (Thermo Fisher). After the first feed on day one, ExpiCHO cells were cultured in a shaker incubator at 33 °C, 115 rpm and 8% CO_2_ following the Max Titer protocol with an additional feed on day five (Thermo Fisher). Culture supernatants were harvested 13 to 14 days after transfection, clarified by centrifugation at 4000 rpm for 20 min, and filtered using a 0.45 μm filter (Thermo Fisher). The AR3A antibody column was used to extract E2mc3-attached nanoparticles from the supernatants, which was followed by SEC on a Superose 6 10/300 GL column. For E2 cores and nanoparticles, protein concentration was determined using UV_280_ absorbance with theoretical extinction coefficients.

### *Blue native* polyacrylamide gel electrophoresis (BN-PAGE)

HCV E2 core nanoparticles were analyzed by *blue native* polyacrylamide gel electrophoresis (*BN*-*PAGE*) and stained with Coomassie blue. The proteins were mixed with sample buffer and G250 loading dye and added to a 4-12% Bis-Tris NativePAGE^TM^ gel (Life Technologies). BN-PAGE gels were run for 2.5 hours at 150 V using the NativePAGE^TM^ running buffer (Life Technologies) according to the manufacturer’s instructions.

### Enzyme-linked immunosorbent assay (ELISA)

Each well of a Costar^TM^ 96-well assay plate (Corning) was first coated with 50 µl PBS containing 0.2 μg of the appropriate antigens. The plates were incubated overnight at 4°C, and then washed five times with wash buffer containing PBS and 0.05% (v/v) Tween 20. Each well was then coated with 150 µl of a blocking buffer consisting of PBS, 40 mg ml^-1^ blotting-grade blocker (Bio-Rad), and 5% (v/v) FBS. The plates were incubated with the blocking buffer for 1 hour at room temperature, and then washed five times with wash buffer. In the mouse sample analysis, serum or plasma was diluted by 50-fold in the blocking buffer and subjected to a 10-fold dilution series. For each sample dilution, a total of 50 μl volume was added to the wells. Each plate was incubated for 1 hour at room temperature, and then washed five times with wash buffer. A 1:2000 dilution of horseradish peroxidase (HRP)-labeled goat anti-mouse IgG antibody (Jackson ImmunoResearch Laboratories) was then made in the wash buffer, with 50 μl of this diluted secondary antibody added to each well. The plates were incubated with the secondary antibody for 1 hour at room temperature, and then washed five times with wash buffer. Finally, the wells were developed with 50 μl of TMB (Life Sciences) for 3-5 min before stopping the reaction with 50 μl of 2 N sulfuric acid. The resulting plate readouts were measured at a wavelength of 450 nm. Of note, the week 2 serum binding did not reach the plateau (or saturation) to allow for accurate determination of EC_50_ titers. Nonetheless, the EC_50_ values calculated in Prism were used as a quantitative measure of antibody titers to facilitate the comparison of different vaccine groups at week 2.

### Bio-layer interferometry (BLI)

The kinetics of E2 cores and nanoparticle binding to HCV-specific antibodies was measured using an Octet Red96 instrument (fortéBio, Pall Life Sciences). All assays were performed with agitation set to 1000 rpm in fortéBio 1× kinetic buffer. The final volume for all the solutions was 200 μl per well. Assays were performed at 30 °C in solid black 96-well plates (Geiger Bio-One). 5 μg ml^-1^ of antibody in 1× kinetic buffer was loaded onto the surface of anti-human Fc Capture Biosensors (AHC) for E2 cores and of anti-human Fc Quantitation Biosensors (AHQ) for nanoparticles for 300 s. A 60 s biosensor baseline step was applied prior to the analysis of the association of the antibody on the biosensor to the antigen in solution for 200 s. A two-fold concentration gradient of antigen, starting at 3.57 μM for E2 cores and 52.08 nM for nanoparticles, depending on the size, was used in a titration series of six. The dissociation of the interaction was followed for 300 s. Correction of baseline drift was performed by subtracting the mean value of shifts recorded for a sensor loaded with antibody but not incubated with antigen and for a sensor without antibody but incubated with antigen. Octet data were processed by FortéBio’s data acquisition software v.8.1. Experimental data were fitted with the binding equations describing a 2:1 interaction to achieve optimal fitting. Of note, E2mc3-v1 binding was also measured using AHQ to facilitate the comparison of antibody binding signals with nanoparticles.

### Differential scanning calorimetry (DSC)

Thermal melting curves of HCV E2 core glycoproteins were obtained with a MicroCal VP-Capillary calorimeter (Malvern). The purified E2 glycoproteins produced from 293S cells were buffer exchanged into 1×PBS and concentrated to 27–50 μM before analysis by the instrument. Melting was probed at a scan rate of 90 °C·h^−1^ from 25 °C to 110 °C. Data processing, including buffer correction, normalization, and baseline subtraction, was conducted using the standardized protocol from the Origin 7.0 software.

### Protein expression and purification for crystallization

The E2 constructs were expressed and purified as previously described (*18, 40*). Fabs AR3A and AR3B were expressed and purified as previously described (*61*). The mAbs were purified on a protein G affinity column followed by size exclusion chromatography using a Superdex-200 column (Pharmacia) in 50 mM NaCl, 20 mM Tris-HCl (pH=7.2) buffer.

### Crystallization and structural determination of HK6a E2c3 – Fab E1 – AR3A – protein G complex

The HK6a E2c3 – Fab E1 – AR3A complex was formed by overnight incubation of purified E2 and Fabs in a molar ratio of 1:1.2:1.25 (E2:Fab E1:Fab AR3A) at room temperature followed by size exclusion chromatography (Superdex-200) to remove unbound Fabs using 20 mM Tris and 50 mM NaCl (pH=7.2) buffer. Crystallization experiments were performed using our high-throughput CrystalMation robotic system (Rigaku) at Scripps Research using the vapor diffusion sitting drop method (drop size 0.3 μl) at 20 °C and resulted in crystals that diffracted to ∼5 Å. To improve crystal resolution, prior to the crystallization experiment, domain III of protein G (PDB entry 1IGC) was added to the HK6a E2c3 – Fab E1 – AR3A complex in a molar ratio of 1:2 (complex: protein G). These experiments resulted in crystals of HK6a E2c3 – Fab E1 – AR3A – protein G that diffracted to 3.40 Å (table S1) using a reservoir solution of 0.2M magnesium chloride, 10% (w/v) PEG 3000, 15% ethylene glycol, 0.1M Na-cacodylate, pH=6.5. Prior to data collection, crystals were flash cooled in liquid nitrogen. Diffraction data sets were collected at Stanford Synchrotron Radiation Lightsource (SSRL) (table S1). Data were integrated and scaled using HKL2000 (*62*) and structure was solved by molecular replacement method using Phaser (*63*) with the HK6a E2c3 - AR3A (PDB entry 6BKB) as a search model. Structure refinement was carried out in Phenix (*64*) and model building with COOT (*65*). Final refinement statistics are summarized in table S1.

### Crystallization and structural determination of E2mc3 - Fab complexes

Crystallization experiments were performed for H77 E2mc3, H77 E2mc3v-1, H77 E2mc3v-6, HK6a E2mc3, and HK6a E2mc3v-1 in complex with AR3A, AR3B, AR3C, and AR3D Fabs. The E2-Fab complexes were formed by overnight incubation of purified E2 and Fabs in a molar ratio of 1:1.25 (E2:Fab) at room temperature followed by size exclusion chromatography (Superdex-200) to remove unbound Fabs using 20 mM Tris and 50 mM NaCl (pH=7.2) buffer. Crystallization screening was again performed on our high-throughput CrystalMation robotic system (Rigaku) using the vapor diffusion sitting drop method (drop size 0.3 μl) at 20 °C and crystals of H77 E2mc3-v1 - AR3C, H77 E2mc3-v6 - AR3C, and HK6a E2mc3-v1 - AR3B were formed that diffracted to 1.90 Å, 2.85 Å, and 2.06 Å, respectively (table S2). Crystals of the H77 E2mc3-v1 - AR3C complex were obtained using a reservoir solution of 20% (w/v) PEG 3500, 0.2M di-ammonium hydrogen phosphate; H77 E2mc3-v6 -AR3C complex from 20% (w/v) PEG 3500, 0.2M Na-thiocyanate, pH=6.9; and HK6a E2mc3-v1 - AR3B complex from 20% (w/v) PEG 8000, 0.1M HEPES pH=7.5. Prior to data collection, H77 E2mc3-v6 - AR3C and HK6a E2mc3-v1 - AR3B crystals were cryoprotected with 10-15% ethylene glycol and flash cooled in liquid nitrogen. Diffraction data sets were collected at the Advanced Photon Source (APS) (table S2). Data were integrated and scaled using HKL2000 (*62*). Structures were solved by molecular replacement method using Phaser (*63*) with the H77 E2c - AR3C or HK6a E2c3 - AR3B (PDB entry 4MWF or 6BKC) as a search model. Structure refinement was carried out in Phenix (*64*) and model building with COOT (*65*). Final refinement statistics are summarized in table S2.

### Negative stain electron microscopy (EM)

The EM experiments were conducted at the Scripps Core Microscopy Facility. Briefly, nanoparticle samples were prepared at the concentration of 0.01 mg/ml. Carbon-coated copper grids (400 mesh) were glow-discharged and 8 µL of each sample was adsorbed for 2 minutes. Excess sample was wicked away and grids were negatively stained with 2% uranyl formate for 2 minutes. Excess stain was wicked away and the grids were allowed to dry. Samples were analyzed at 80kV with a Talos L120C transmission electron microscope (Thermo Fisher) and images were acquired with a CETA 16M CMOS camera.

### Mouse immunization and sample collection

The Institutional Animal Care and Use Committee (IACUC) guidelines were followed with animal subjects tested in the immunization study. Eight-week-old BALB/c mice were purchased from The Jackson Laboratory. Mice were housed in ventilated cages in environmentally controlled rooms at Scripps Research, in compliance with an approved IACUC protocol and AAALAC guidelines. Mice were immunized at weeks 0, 3, 6 and 9 for a total of four times. Each immunization consisted of 200 μl of antigen/adjuvant mix containing 50 μg of vaccine antigen and 100 μl of AddaVax adjuvant (Invivogen) via the subcutaneous (s.c.) route. Blood was collected two weeks after each immunization. All bleeds were performed through the facial vein (submandibular bleeding) using lancets (Goldenrod). While intermediate bleeds were collected without anticoagulant, terminal bleeds were collected using EDTA-coated tubes. Serum and plasma were heat inactivated at 56 °C for 30 min, spun at 1000 RPM for 10 min, and sterile filtered. The cells were washed once in PBS and then resuspended in 1 ml of ACK Red Blood Cell lysis buffer (Lonza). After two rounds of washing with PBS, peripheral blood mononuclear cells (PBMCs) were resuspended in 2 ml of Bambanker Freezing Media (Lymphotec). In addition, spleens were also harvested and grounded against a 70-μm cell strainer (BD Falcon) to release the splenocytes into a cell suspension. Splenocytes were centrifuged, washed in PBS, treated with 5 ml of Red Blood Cell Lysis Buffer Hybri-Max (Sigma-Aldrich), and frozen with 10% of DMSO in FBS. While serum and plasma were used in HCV neutralization assays, 80% of the plasma from individual mice at week 11 in study #1 (9, 10, and 10 in groups 1, 2, and 3, respectively) was purified using a 0.2-ml protein G spin kit (Thermo Scientific) following the manufacturer’s instructions. Purified IgGs were used to assess the polyclonal NAb response in HCV neutralization assays.

### HCV neutralization assay

HCV pseudotyped particle (HCVpp) assays were utilized to assess the neutralizing activity of vaccine-induced antibody response in mouse sera, as well as synthesized antibodies from the next-generation sequencing (NGS) analysis of bulk-sorted mouse splenic B cells. Briefly, HCVpps were generated by co-transfection of 293T cells with pNL4-3.lucR-E-plasmid and the corresponding expression plasmids encoding the E1E2 genes at a 4:1 ratio by polyethylenimine as previously described (*66*). In vitro neutralization was carried on Huh7.5 cells using a single dilution of 1:50 for mouse sera and three concentrations (10μg/ml, 1.0μg/ml, and 0.1μg/ml) for antibodies. Full neutralization curves were determined for IgGs purified from mice in study #1 against autologous H77 (1a) and heterologous SA13 (5a), with a starting IgG concentration of 100 μg/ml and a series of three-fold dilutions.

### Bulk sorting of HCV E2-specific mouse B cells

Spleens were harvested from immunized mice 15 days after the last immunization and cell suspension was prepared. Cells were stained as follows: dead cells were excluded by staining with Fixable Aqua Dead Cell Stain kit (Thermo Fisher L34957). Receptors FcγIII (CD16) and FcγII (CD32) were blocked by adding 20 µl of 2.4G2 mAb (BD Pharmigen N553142). Cells were then incubated with 10 μg/ml of biotinylated HCV E2mc3 protein. Briefly, E2mc3 was generated by biotinylation of the individual Avi-tagged HCV E2mc3 using biotin ligase BirA according to the manufacturer’s instructions (Avidity LLC). Biotin excess was removed by SEC on a Superdex 200 column (GE Healthcare). In the SEC profile, the Avi-tagged E2mc3 peak is centered at 14.5 ml, while a broader peak of biotin ligase can be found at 18-23 ml. Cells and biotinylated proteins were incubated for 5 min at 4 °C, followed by the addition of 2.5 µl of anti-mouse IgG fluorescently labeled with FITC (Jackson ImmunoResearch 115-095-071) and incubated for 15 min at 4 °C. Finally, 5 µl of premium-grade allophycocyanin (APC)-labeled streptavidin were added to the cells and incubated for 15 min at 4 °C. In each step, cells were washed with DPBS and the sorting buffer was 0.5 ml FACS buffer. FITC^+^ APC^+^ E2mc3 specific B cells were sorted using BD FACSAria II into Eppendorf tube with 500 μl of FACS buffer.

### Next-generation sequencing (NGS) and bioinformatics analysis of mouse B cells

A 5’-rapid amplification of cDNA ends (RACE) protocol has been reported for unbiased sequencing of mouse B cell repertoires (*26, 29*). Here, this protocol was applied to bulk-sorted, E2-specific mouse splenic B cells. Briefly, 5’-RACE cDNA was obtained from bulk-sorted splenic B cells of each mouse with SMART-Seq v4 Ultra Low Input RNA Kit for Sequencing (TaKaRa). The immunoglobulin PCRs were set up with Platinum *Taq* High-Fidelity DNA Polymerase (Life Technologies) in a total volume of 50 µl, with 5 μl of cDNA as template, 1 μl of 5’-RACE primer, and 1 μl of 10 µM reverse primer. The 5’-RACE primer contained a PGM/S5 P1 adaptor, while the reverse primer contained a PGM/S5 A adaptor. We adapted the mouse 3’-Cγ1-3/3’-Cμ inner primers and 3’-mCκ outer primer as reverse primers for 5’-RACE PCR processing of heavy and light (κ) chains. A total of 25 cycles of PCR was performed and the expected PCR products (500- 600 bp) were gel purified (Qiagen). NGS was performed on the Ion S5 GeneStudio system. Briefly, heavy and light (κ) chain libraries from the same mouse were quantitated using Qubit® 2.0 Fluorometer with Qubit® dsDNA HS Assay Kit, and then mixed using a ratio of 3:1 before being pooled with antibody libraries of other mice at an equal ratio for sequencing. Template preparation and (Ion 530) chip loading were performed on Ion Chef using the Ion 520/530 Ext Kit, followed by sequencing on the Ion S5 system with default settings. The mouse *Antibodyomics* pipeline (*29*) was used to process the raw data and to determine distributions for germline gene usage, somatic hypermutation (SHM), germline divergence, and H/KCDR3 loop length.

## List of Supplementary Online Materials (SOM)

**Fig. S1.**
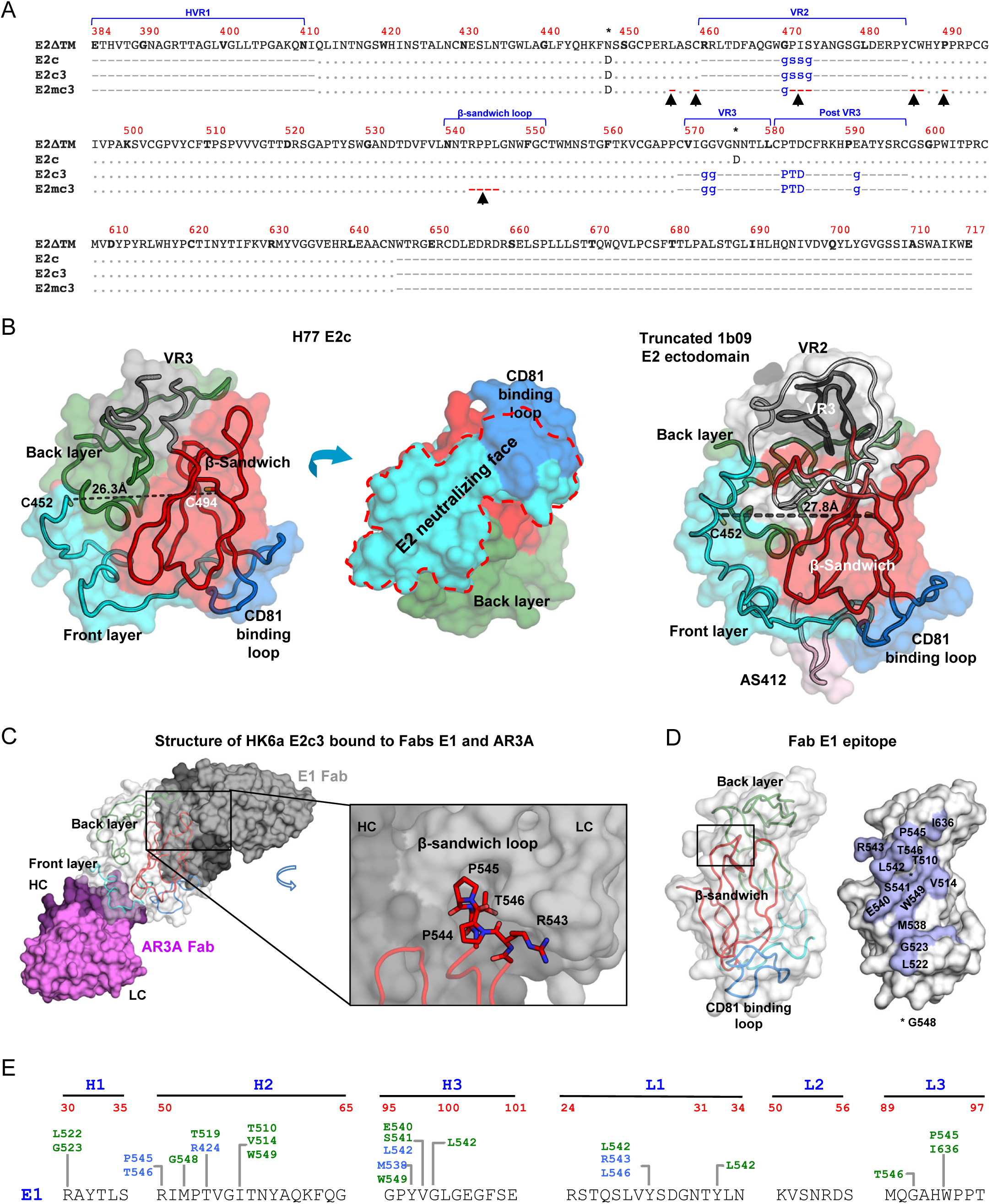
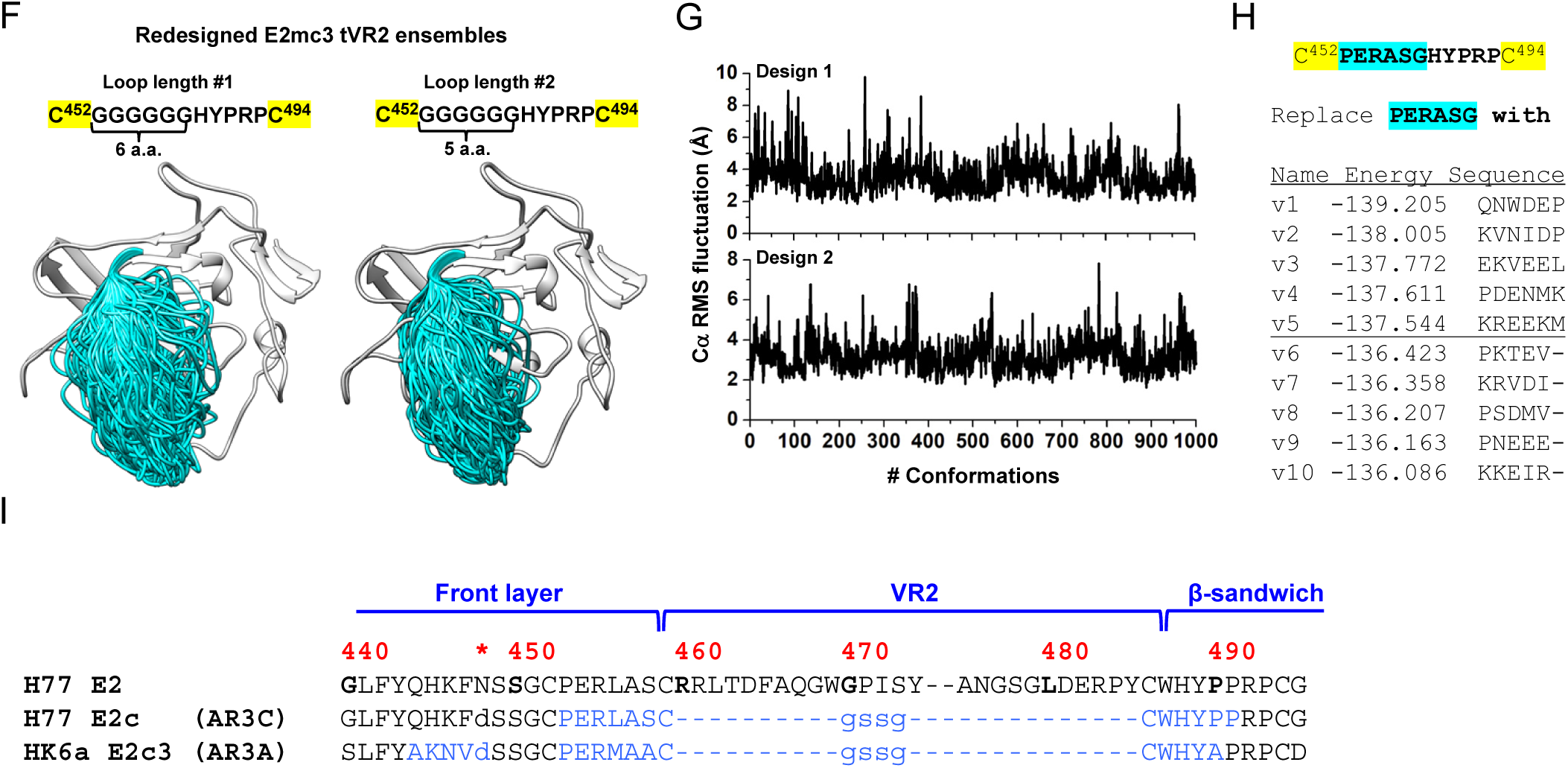
Sequence, structural, and computational analyses of HCV envelope glycoprotein E2 and E2 core design variants. **(A)** Sequence alignment of H77 E2ΔTM, E2c, E2c3, and E2mc3 constructs. Regions of HVR1, VR2, VR3, and the β-sandwich loop are marked with blue lines. The E2c and E2c3 mutations are labeled in blue and the E2mc3 mutations in red and with arrows. **(B)** Structures of H77 E2c (PDB: 4MWF) and 1b09 truncated E2 ectodomain (PDB: 6MEI). The protein chain is represented as a tube with the molecular surface color-coded as in Fig. 1A. **(C)** Structure of HK6a E2c3 bound to Fabs E1 and AR3A. The molecular surfaces of HK6a E2c3, E1, and AR3A are shown in light gray, dark gray, and magenta, respectively. A close-up view of four Fab E1-interacting amino acids at the tip of the β-sandwich loop is shown as an insert. **(D)** Projection of the Fab E1 epitope onto the E2c3 structure. Left: back layer, β-sandwich loop, and CD81 binding loop are shown as tubes within the transparent gray molecular surface. The tip of the β-sandwich loop is labeled with a rectangle; Right: Fab E1-interacting amino acids are labeled on the solid molecular surface of E2 that is colored in cornflower blue. **(E)** Schematic overview of the interactions between E1 mAb HC and LC CDR and E2. E2 interacting residues are highlighted in blue (hydrogen bonds) and green (hydrophobic contacts). **(F)** Conformational ensembles of redesigned E2mc3 tVR2 loop. E2mc3 structure is shown in gray ribbons and 1000 modeled loops are shown in cyan tubes. Left: loop length #1 (13 a.a.); right: loop length #2 (12 a.a.). **(G)** Distribution of Cα root-mean-square (RMS) fluctuation plotted for the two redesigned tVR2 loop ensembles. **(H)** Five top-ranking designs and their energy scores for the two loop ensembles. **(I)** Sequence alignment of H77 E2, H77 E2c, and HK6a E2c3. Amino acids that are disordered in the crystal structures of H77 E2c-AR3C (PDB: 4MWF) and HK6a E2c3-AR3A (PDB: 6BKB) complexes are colored in light blue.

**Fig. S2.**
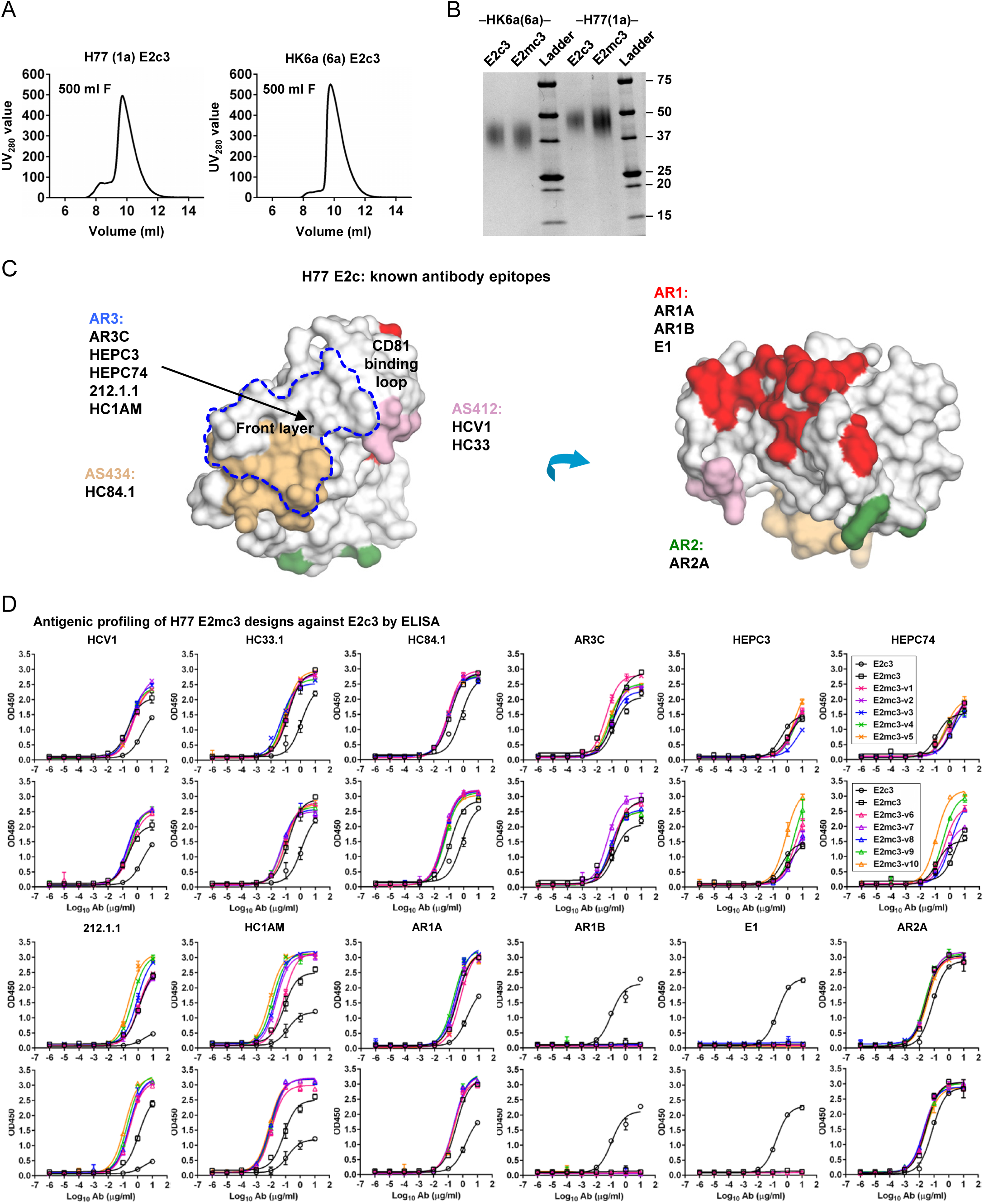
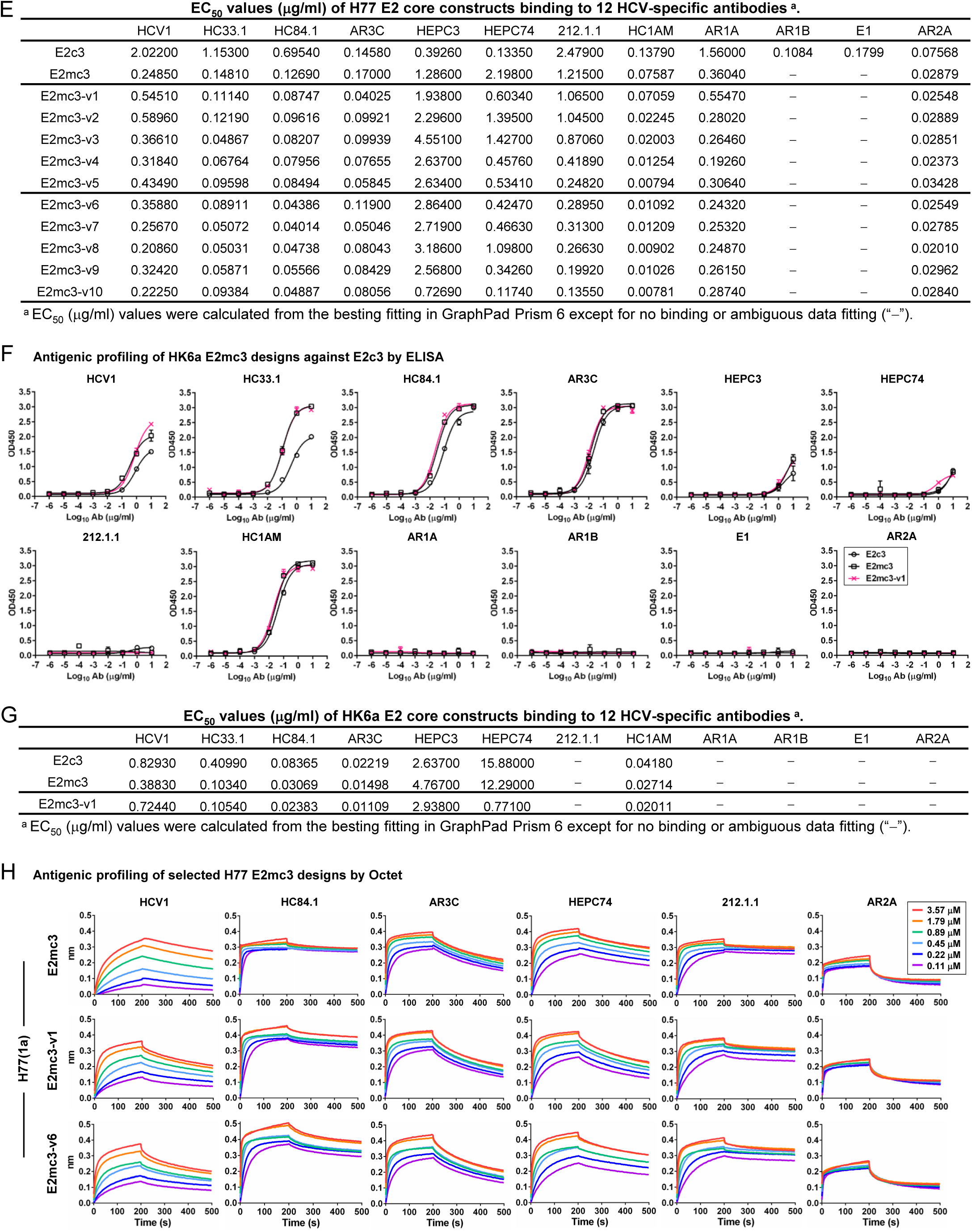
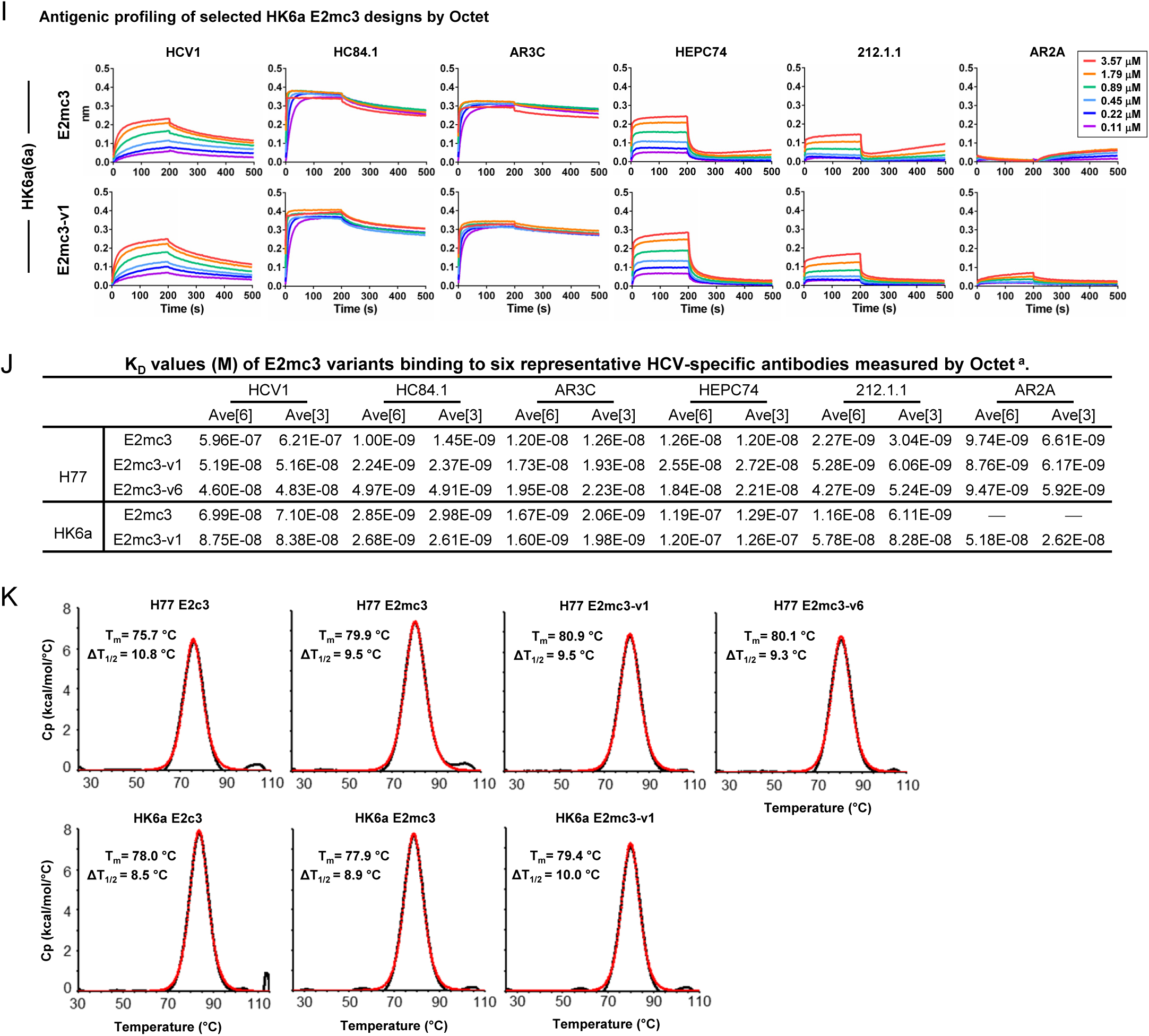
Biochemical, biophysical, and antigenic characterization of E2 cores derived from H77(1a) and HK6a(6a). (A) SEC profiles of E2c3 proteins obtained from a Superdex 200 10/300 column after immunoaffinity (AR3A) purification. (B) SDS-page of E2c3 and E2mc3 proteins after immunoaffinity (AR3A) and SEC purification. (**C**) Antigenic sites and epitopes mapped onto the H77 E2c surface using the following mAbs panel: AR3C (18, 43), HEPC3/74 (44), 212.1.1 (47), HC1AM (68), HC84.1 (69), HCV1 (70), HC33 (71), AR1A/B and AR2A (43, 72), and E1 (61). (**D**) ELISA binding of H77 E2c3 and E2mc3 variants to 12 HCV-specific antibodies. **(E)** EC_50_ values (μg/ml) of H77 E2 core constructs binding to 12 HCV-specific antibodies. (**F**) ELISA binding of HK6a E2 E2c3 and E2mc3 variants to 12 HCV-specific antibodies. (**G**) EC_50_ values (μg/ml) of HK6a E2 core constructs binding to 12 HCV-specific antibodies. In (**E**) and (**G**), EC_50_ values were calculated for all ELISA plots in Prism except where the highest OD_450_ value was below 0.1 or data fitting was ambiguous (denoted as “−”). (**H**) Octet binding of H77 E2mc3 variants to six HCV-specific antibodies. (**I**) Octet binding of HK6a E2mc3 variants to six HCV-specific antibodies. In (**H**) and (**I**), sensorgrams were obtained from an Octet RED96 instrument using a titration series of six concentrations (3.57-0.11 μM by twofold dilution for all E2mc3 variants) and kinetics biosensors (see Methods). (**J**) K_D_ (M) values of H77 and HK6a E2mc3 variants measured by Octet. **(K)** Differential scanning calorimetry (DSC) curves of selected E2 core constructs. Two thermal parameters, T_m_ and T_1/2_, are labeled on the DSC profiles.

**Fig. S3.**
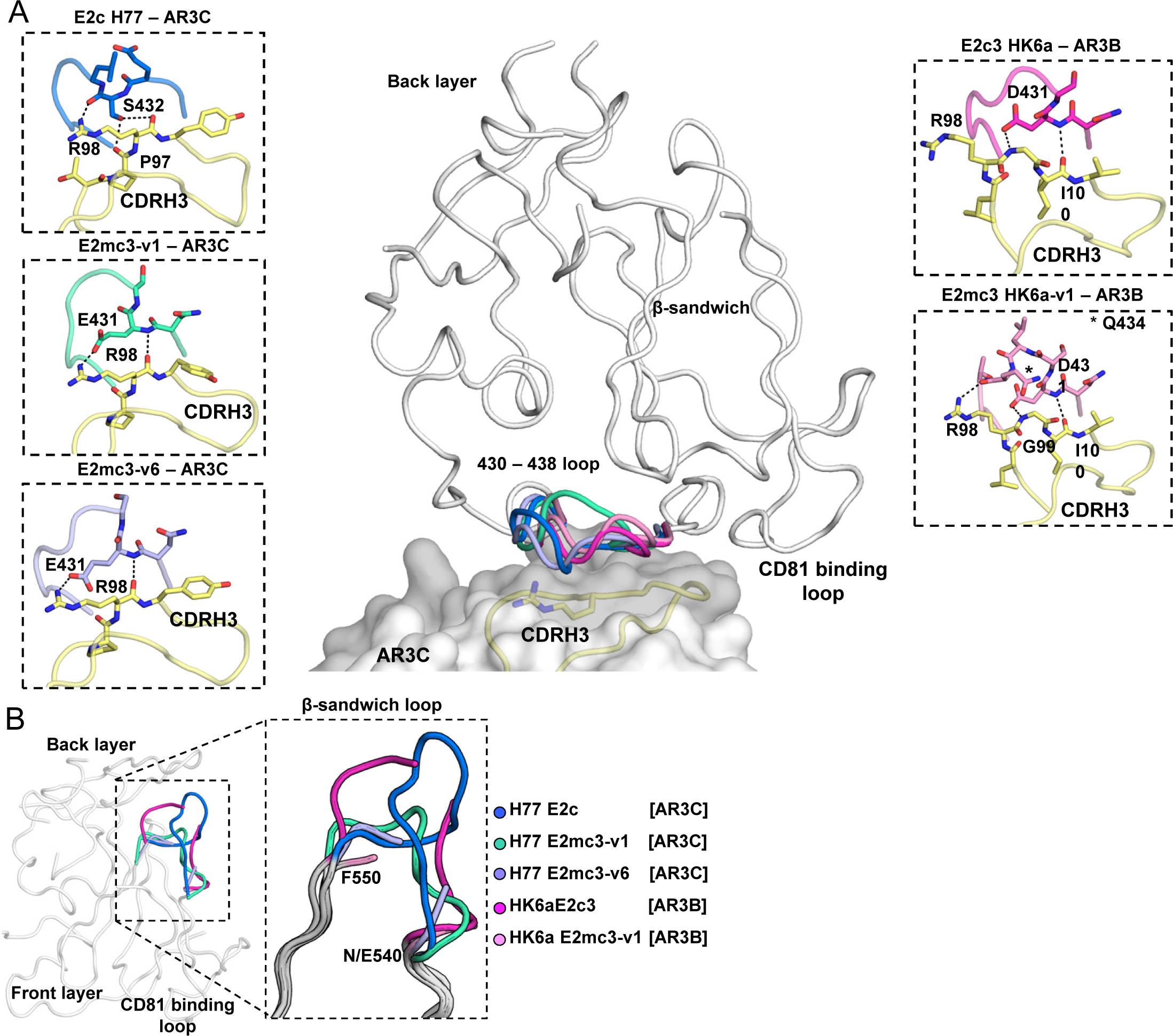
Crystal structures of H77 E2mc3 and HK6a E2mc3. (**A**) Conformational flexibility of the front layer 430-438 loop. The 430-438 loop in the H77 and HK6a E2c structures acquire different conformations yet maintain similar interactions with the Fab CDRH3 loop, indicating high flexibility of this region. (**B**) The conformation of the β-sandwich loop (a.a. 538-553) in H77 E2c, HK6a E2c3, and H77/HK6a E2mc3.

**Fig. S4.**
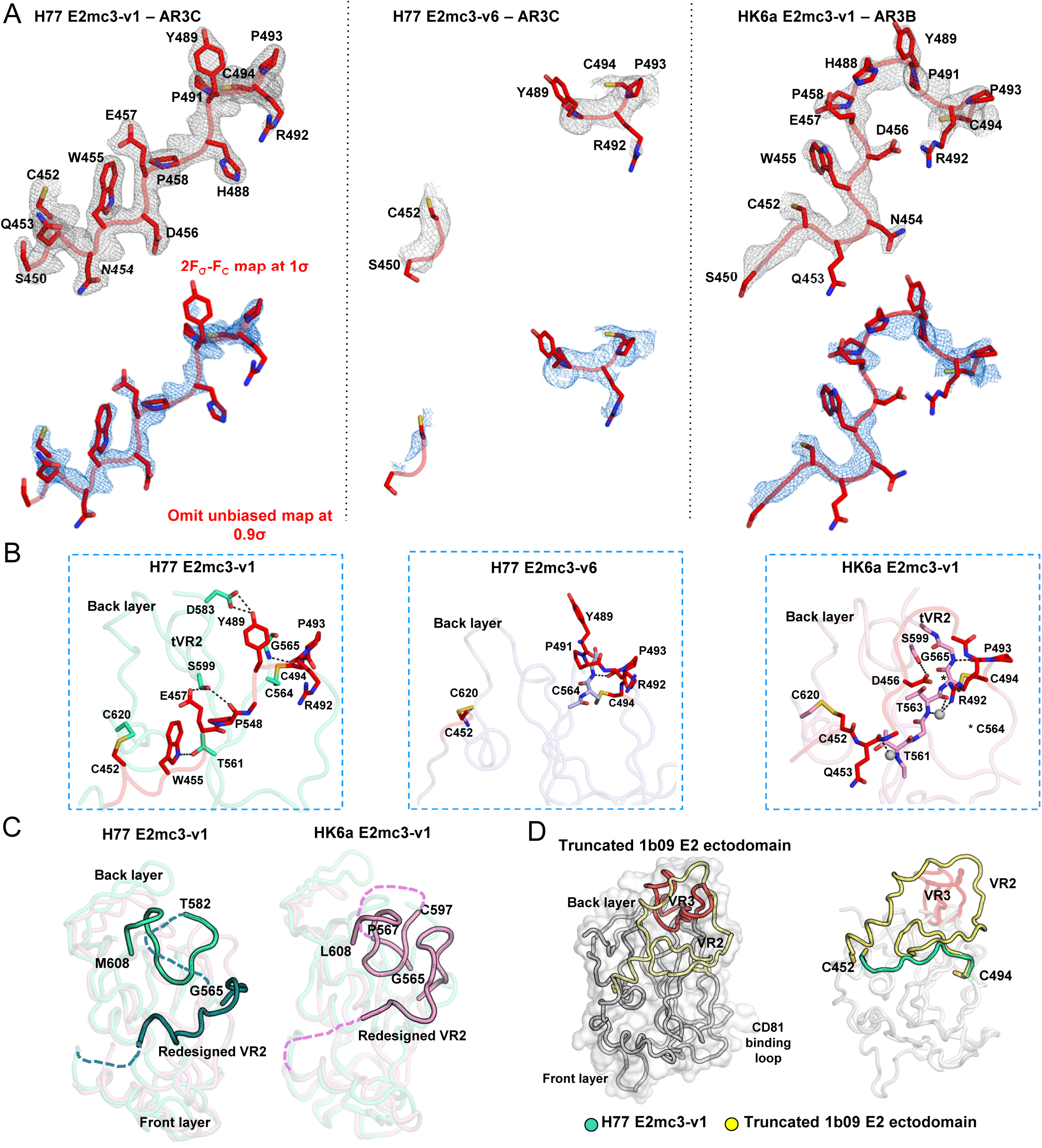
Structural analysis of the redesigned tVR2 region. (**A**) 2F_O_-F_C_ and unbiased omit electron density maps of the redesigned VR2 region (1σ or 0.9σ, respectively). (**B**) The intrinsic interactions of the redesigned tVR2 region of H77 E2mc3-v1, H77 E2mc3-v6, and HK6a E2mc3-v1 with the VR3 region (a.a. 570-597). (**C**) Comparison of the H77 E2mc3-v1 and HK6a E2mc3-v1 indicating conformational changes of the redesigned VR2 and the neighboring VR3-back layer region (a.a. 565-608). (**D**) In the crystal structure of the truncated 1b09 E2 ectodomain (left), the full length VR2 wraps around VR3 (colored by yellow and red) to form the variable face. Superposition of H77 E2mc3-v1 structure on 1b09 E2 indicates only a minor influence of the VR2 redesign on the E2 overall fold (as indicated by aligning C452 and C494).

**Fig. S5.**
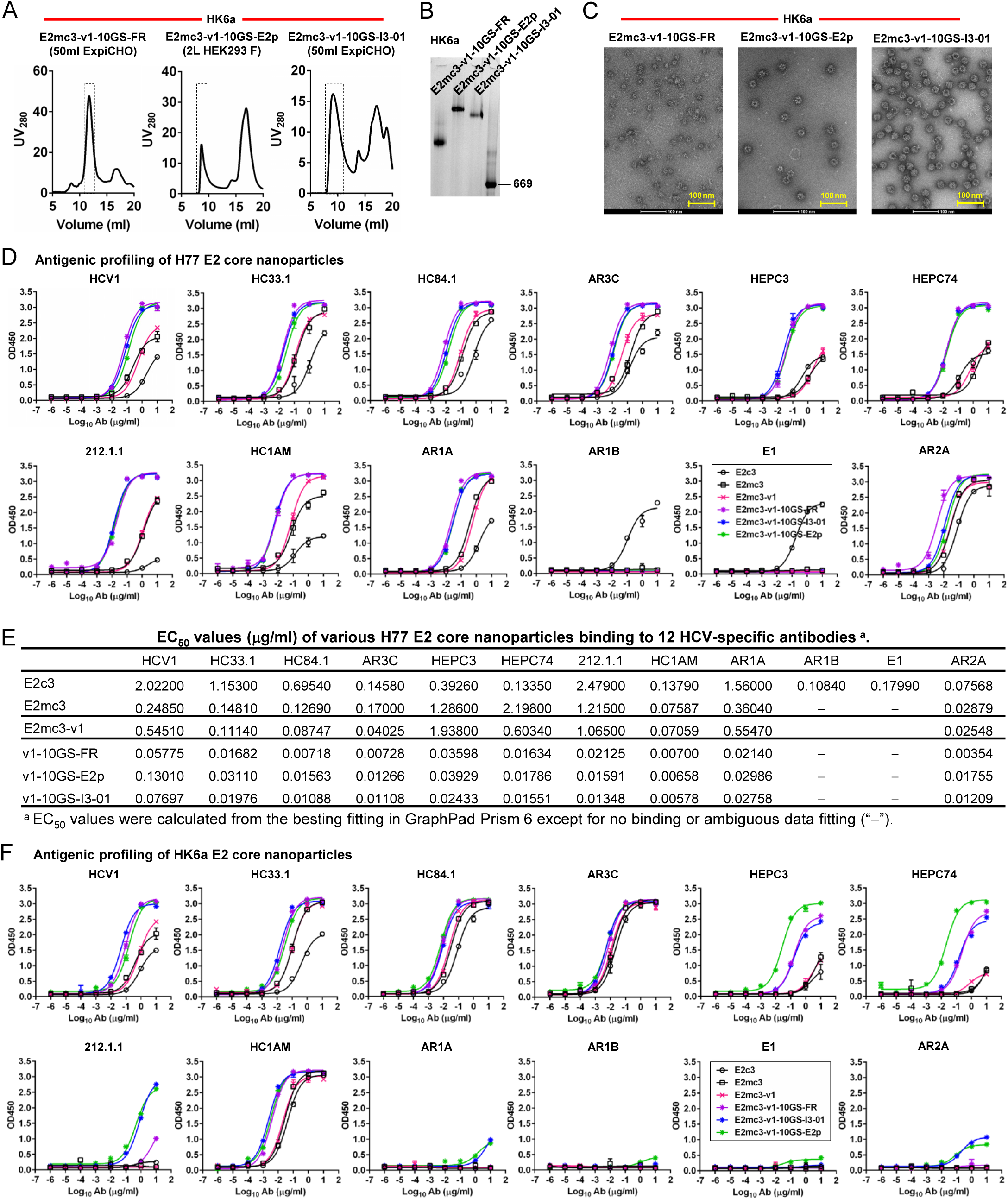
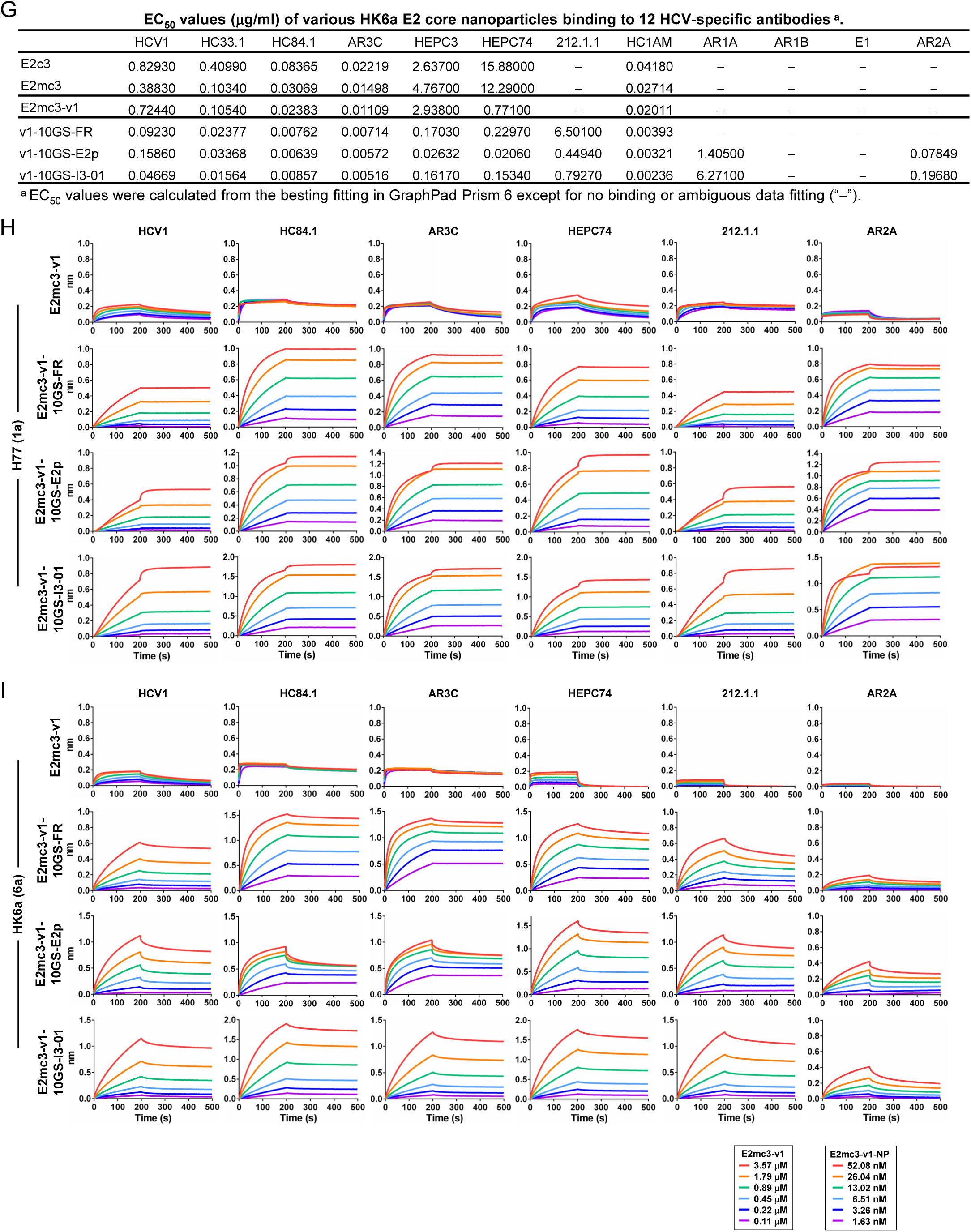
Biochemical, biophysical, and antigenic characterization of E2 core nanoparticles derived from H77(1a) and HK6a(6a). (**A**) SEC profiles of HK6a E2mc3-v1 nanoparticles based on FR, E2p and I3-01 obtained from a Superose 6 200 Increase 10/300 column after immunoaffinity (AR3A) purification. (**B**) BN-PAGE of HK6a E2mc3-v1 nanoparticles based on FR, E2p and I3-01. (**C**) Negative-stain EM images of HK6a E2mc3-v1 nanoparticles based on FR, E2p and I3-01. (**D**) ELISA binding of H77 E2c3-v1 nanoparticles to 12 HCV-specific antibodies. (**E**) EC_50_ values (μg/ml) of H77 E2 E2c3-v1 nanoparticles binding to 12 HCV-specific antibodies. (**F**) ELISA binding of HK6a E2mc3-v1 nanoparticles to 12 HCV-specific antibodies. (**G**) EC_50_ values (μg/ml) of HK6a E2mc3-v1 nanoparticles with 12 HCV-specific antibodies. In (**E**) and (**G**), EC_50_ values were calculated for all ELISA plots in GraphPad Prism 6 except where the highest OD_450_ value was below 0.1 or data fitting was ambiguous (denoted as “−”). (**H**) Octet binding of H77 E2mc3-v1 nanoparticles to six HCV-specific antibodies. (**I**) Octet binding of HK6a E2mc3-v1 nanoparticles to six HCV-specific antibodies. In (**H**) and (**I**), sensorgrams were obtained from an Octet RED96 instrument using a titration series of six concentrations (3.57-0.11 μM by twofold dilution for E2mc3-v1 and 52.08-1.63 nM by twofold dilution for E2mc3-v1 nanoparticles) and quantitation biosensors (see Materials and Methods).

**Fig. S6.**
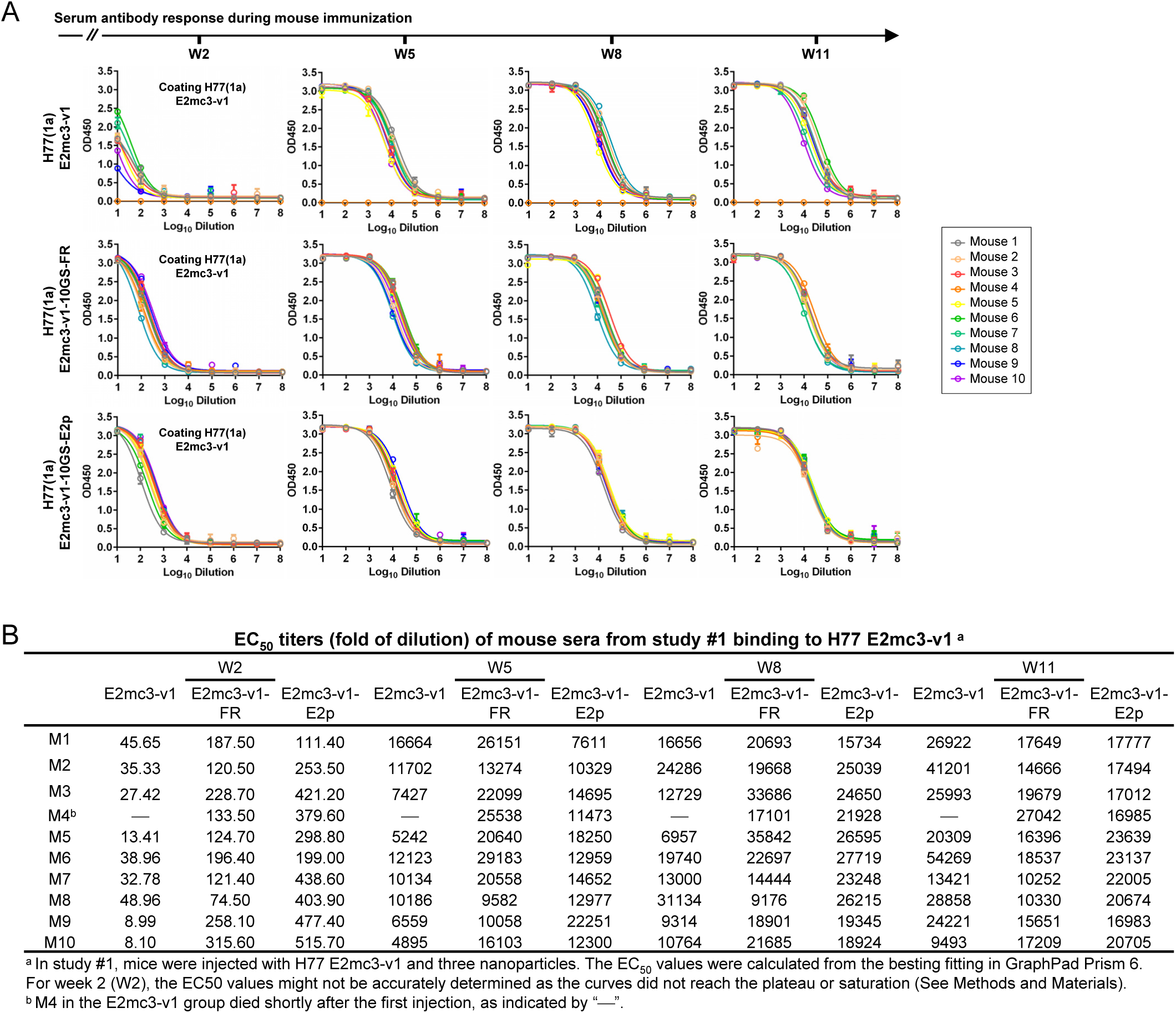
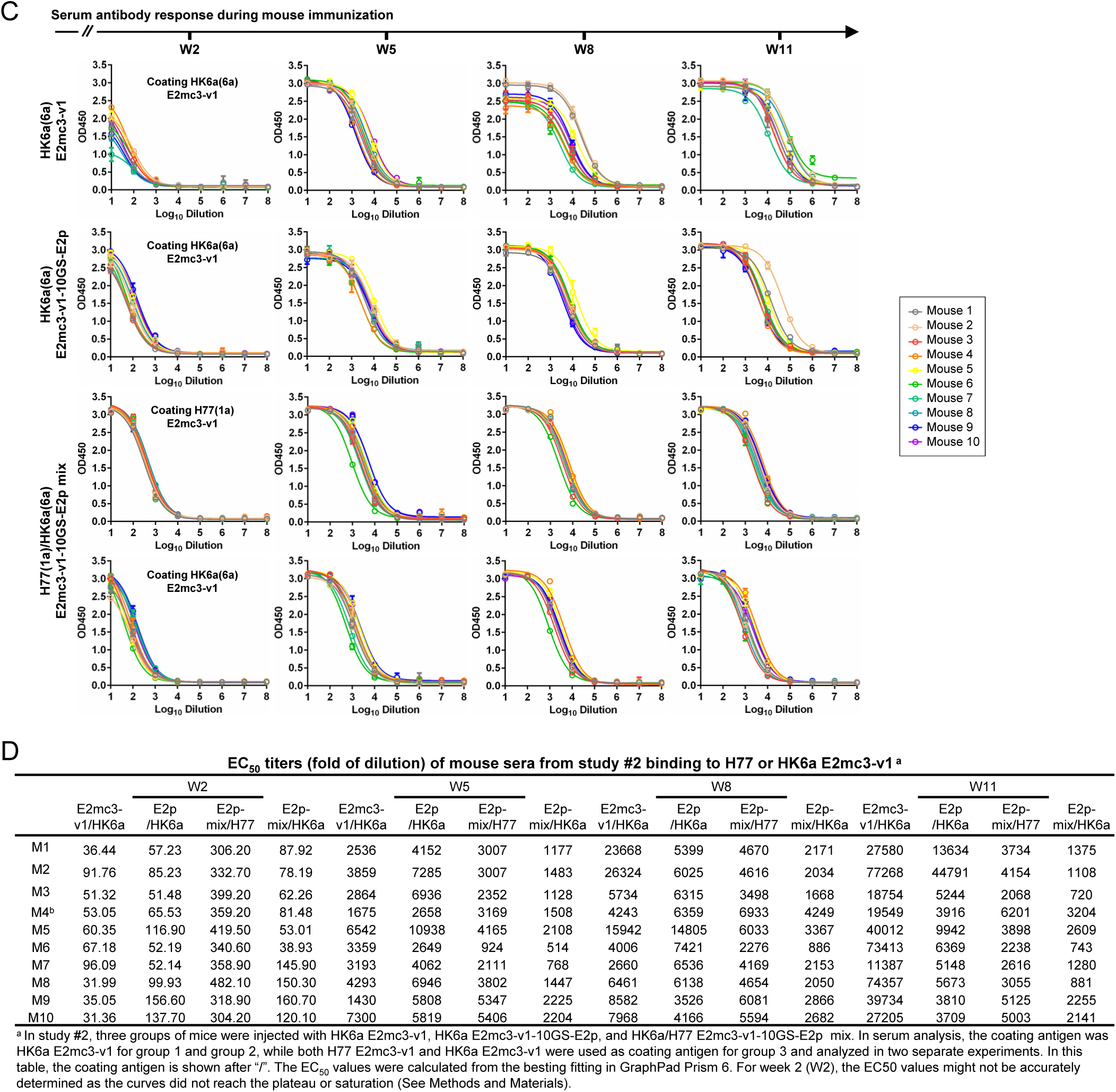
Murine antibody response during immunization at w2, w5, w8 and w11. (**A**) ELISA binding of H77 E2mc3-v1 to mouse sera from groups 1, 2, and 3 in study #1, which were immunized with H77 E2mc3-v1, E2mc3-v1-10GS-FR, and E2mc3-v1-10GS-E2p, respectively, at four time points. (**B**) EC_50_ titers (fold of dilution) of study #1 mouse sera binding to H77 E2mc3-v1 at four time points. (**C**) ELISA binding of HK6a E2mc3-v1 or H77 E2mc3-v1 to mouse sera from groups 1, 2, and 3 in study #2, which were immunized with HK6a E2mc3-v1, HK6a E2mc3-v1-10GS-E2p, and H77/HK6a E2mc3-v1-10GS-E2p mix, respectively, at four time points. Panels 1 and 2: sera from mice immunized with HK6a E2mc3-v1 (group 1) and HK6a E2mc3-v1-10GS-E2p (group 2) were tested against HK6a E2mc3-v1. Panels 3 and 4: sera from mice immunized with H77/HK6a E2mc3-v1-10GS-E2p mix (group 3) were tested against H77 E2mc3-v1 (panel 3) and HK6a E2mc3-v1 (panel 4). (**D**) EC_50_ titers (fold of dilution) of study #2 mouse sera binding to HK6a or H77 E2mc3-v1 at four time points.

**Fig. S7.**
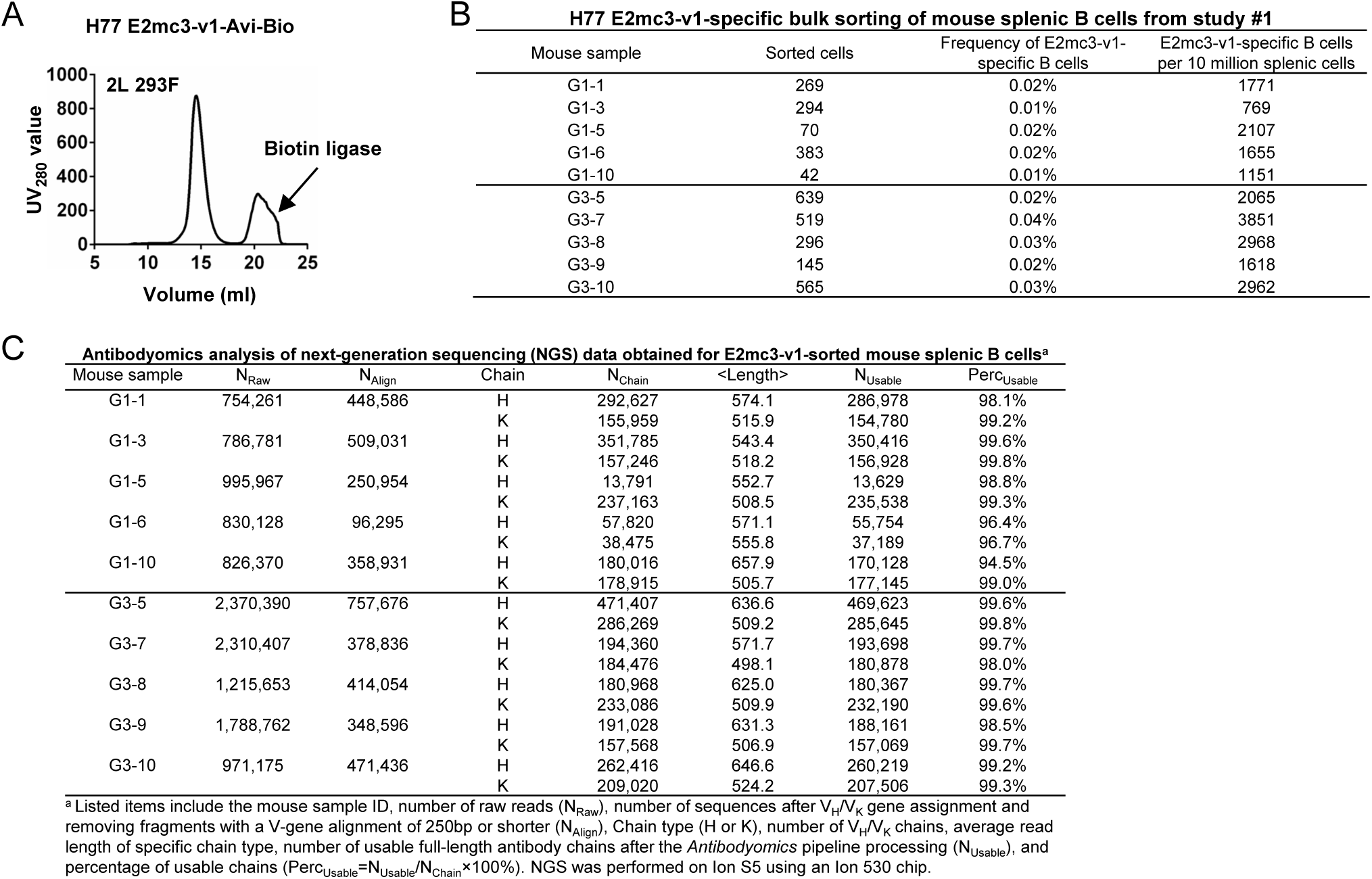
Next-generation sequencing (NGS) analysis of bulk-sorted E2mc3-specific mouse splenic B cells. (**A**) SEC profile of biotinylated Avi-tagged H77 E2mc3-v1, termed E2mc3-v1-Avi-Biot, obtained from a Superdex 200 10/300 column, with the peak corresponding to biotin ligase labeled on the profile. (**B**) Summary of H77 E2mc3-v1-specific bulk sorting of mouse splenic B cells from study #1, groups 1 and 3. (**C**) Antibodyomics analysis of NGS data obtained for E2mc3-v1-sorted mouse splenic B cells. NGS data from groups 1 and 3, a total of 10 mice, were analyzed.

**Fig. S8.**
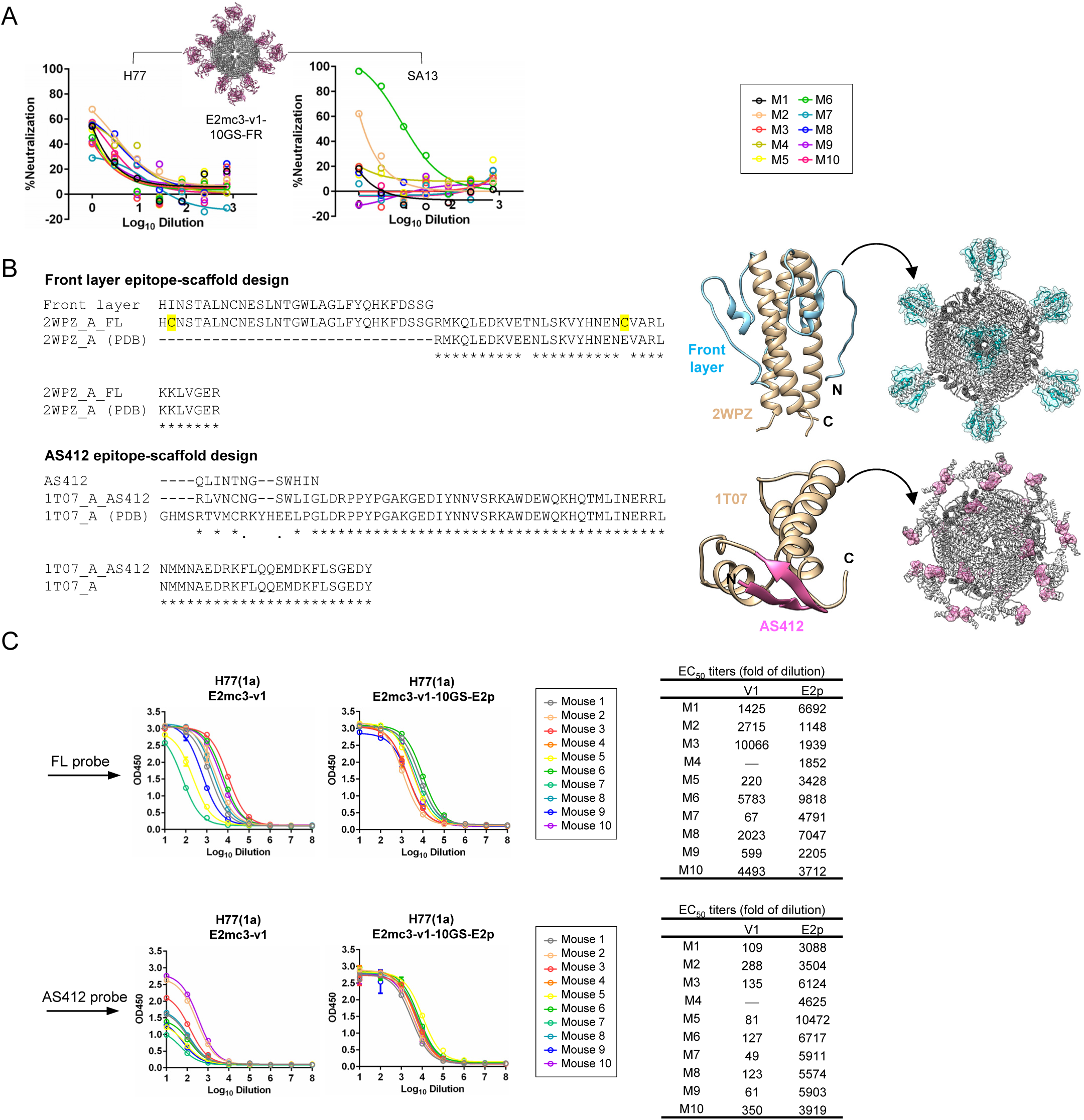
Analysis of mouse polyclonal serum antibody response. (**A**) Neutralization of H77 and SA13 by mouse IgG purified from the H77 E2mc3-v1-10GS-FR group in study #1. The HCVpp neutralization assays were performed with a starting IgG concentration of 100μg/ml and a series of 3-fold dilutions. The full neutralization curves were created to facilitate the comparison between different groups. (**B**) Design of epitope-specific probes for front layer (FL) and AS412. Left: epitope-scaffold design showing sequence alignment of the epitope, the designed epitope-scaffold, and the original scaffold obtained from the database search (* – match and • – similar with engineered disulfide bonds in yellow); Middle: structural model of designed epitope-scaffold, with the scaffold backbone shown in tan and the FL and AS412 epitopes in cyan and pink, respectively. Right: molecular model of nanoparticle probe, with FL and AS412 epitopes shown in cyan and pink, respectively. **(C)** ELISA binding of mouse sera from groups 1 and 3 in study #1, which were immunized with H77 E2mc3-v1 and E2mc3-v1-10GS-E2p, respectively, to the two epitope probes. Left: ELISA curves; Right: summary of EC_50_ titers (fold of dilution) in serum binding analysis.

**Table S1.** Data collection and refinement statistics for HK6a E2c3–Fab E1–Fab AR3A complex structures.

| Data collection | HK6a E2c3 Fab E1 Fab AR3A protein G |
| --- | --- |
| Beamline | SSRL 12-2 |
| Wavelength (Å) | 0.97946 |
| Space Group | P6<sub>3</sub>22 |
| Unit cell parameters (Å, °) | a=b=209.8, c=139.9, γ=120 |
| Resolution range (Å) | 30.00-3.40 (3.46-3.40)<sup>a</sup> |
| Observations | 445,765 |
| Unique reflections | 25,916(1,258) |
| Completeness (%) | 100.0 (99.8) |
| I/σ(I) | 13.5 (1.1) |
| CC<sub>1/2</sub><sup>b</sup> | 0.92 (0.29) |
| R<sub>sym</sub><sup>c</sup> | 0.23 (1.61) |
| R<sub>pim</sub><sup>d</sup> | 0.05 (0.53) |
| Redundancy | 17.2 (9.9) |
| **Refinement Statistics** |  |
| Resolution (Å) | 30.00-3.40 (3.46-3.40) |
| No. reflections | 25,879 |
| R<sub>cryst</sub><sup>e</sup>/R<sub>free</sub><sup>f</sup> | 0.255/0.308 |
| No. atoms |  |
| Protein | 7,680 |
| Glycan | 56 |
| Wilson B (Å<sup>2</sup>) | 100 |
| Average B value (Å<sup>2</sup>) |  |
| All proteins | 101 |
| HK6a E2c3 | 82 |
| Fab E1 | 109 |
| Fab AR3A | 96 |
| protein G | 127 |
| All glycans | 117 |
| R.m.s. deviation from ideal geometry |  |
| Bond length (Å) | 0.002 |
| Bond angle (°) | 0.5 |
| Ramachandran Plot (%)<sup>g</sup> |  |
| Favored | 95.0 |
| Outliers | 0.3 |
| PDB ID | DDDD |

<sup>a</sup> Parentheses refer to outer shell statistics.<sup>b</sup> CC<sub>1/2</sub> = Pearson Correlation Coefficient between two random half datasets.<sup>c</sup> $R_{\text{sym}} = \frac{\sum_{hkl} \sum_i |I_{hkl,i} - \langle I_{hkl} \rangle|}{\sum_{hkl} \sum_i I_{hkl,i}}$ , where $I_{hkl,i}$ is the scaled intensity of the $i^{\text{th}}$ measurement of reflection $h, k, l$ , and $\langle I_{hkl} \rangle$ is the average intensity for that reflection.<sup>d</sup> $R_{\text{pim}} = \frac{\sum_{hkl} (1/(n-1))^{1/2} \sum_i |I_{hkl,i} - \langle I_{hkl} \rangle|}{\sum_{hkl} \sum_i I_{hkl,i}}$ , where $n$ is the redundancy.<sup>e</sup> $R_{\text{cryst}} = \frac{\sum_{hkl} |F_o - F_c|}{\sum_{hkl} |F_o|} \times 100$ , where $F_o$ and $F_c$ are the observed and calculated structure factors.<sup>f</sup> $R_{\text{free}}$ was calculated as for $R_{\text{cryst}}$ , but on a test set of 5% of the data excluded from refinement.<sup>g</sup> Calculated using MolProbity (67)

**Table S2.** Data collection and refinement statistics for H77 and HK6a E2mc3–Fab complex structures.

| Data collection | H77 E2mc3-v1/Fab<br>AR3C | H77 E2mc3-v6/Fab<br>AR3C | HK6a E2mc3-v1/Fab<br>AR3B |
| --- | --- | --- | --- |
| Beamline | APS-23 IDB | APS-23 IDD | APS-23 IDB |
| Wavelength (Å) | 1.0332 | 1.0332 | 1.0332 |
| Space Group | P2<sub>1</sub> | P1 | C2 |
| Unit cell parameters<br>(Å, °) | a=88.4, b=93.6, c=96.7,<br>β=100.4 | a=45.1, b=90.8, c=94.0,<br>α=84.5, β=78.1, γ=77.0 | a=175.2, b=54.0, c=71.4,<br>β=94.7 |
| Resolution range (Å) | 30.00-1.90 (1.93-1.90)<sup>a</sup> | 30.00-2.85 (2.90-2.85) | 50.00-2.06 (2.10-2.06) |
| Observations | 522,659 | 107,943 | 180,390 |
| Unique reflections | 121,242 (5,954) | 29,145 (836) | 38,846 (1,408) |
| Completeness (%) | 99.0 (98.3) | 88.5 (51.3) | 93.9 (68.6) |
| I/σ(I) | 25.8 (1.2) | 14.2 (2.4) | 15.9 (3.5) |
| CC<sub>1/2</sub><sup>b</sup> | 0.89 (0.50) | 0.96 (0.88) | 0.96 (0.92) |
| R<sub>sym</sub><sup>c</sup> | 0.05 (1.16) | 0.09 (0.31) | 0.09 (0.24) |
| R<sub>pim</sub><sup>d</sup> | 0.03 (0.61) | 0.05 (0.21) | 0.04 (0.14) |
| Redundancy | 4.5 (4.3) | 3.7 (2.3) | 4.6 (3.1) |
| **Refinement Statistics** |  |  |  |
| Resolution (Å) | 30.00-1.90 (1.93-1.90) | 30.00-2.85 (2.90-2.85) | 50.00-2.06 (2.10-2.06) |
| No. reflections | 121,189 | 29,122 | 38,835 |
| R<sub>cryst</sub><sup>e</sup>/R<sub>free</sub><sup>f</sup> | 0.207/0.243 | 0.216/0.261 | 0.183/0.228 |
| No. atoms |  |  |  |
| Protein | 8893 | 8748 | 4479 |
| Glycan | 112 | 126 | 120 |
| Water | 520 | 0 | 307 |
| Wilson B (Å<sup>2</sup>) | 35 | 61 | 29 |
| Average B value (Å<sup>2</sup>) |  |  |  |
| All proteins | 47 | 67 | 35 |
| E2mc3 | 44 | 72 | 41 |
| Fab | 48 | 64 | 33 |
| All glycans | 80 | 103 | 63 |
| Water | 47 | - | 38 |
| **R.m.s. deviation from ideal geometry** |  |  |  |
| Bond length (Å) | 0.008 | 0.002 | 0.004 |
| Bond angle (°) | 0.9 | 0.6 | 0.7 |
| **Ramachandran Plot (%)<sup>g</sup>** |  |  |  |
| Favored | 96.8 | 95.9 | 97.4 |
| Outliers | 0.0 | 0.2 | 0.1 |
| PDB ID | AAAA | BBBB | CCCC |

<sup>a</sup> Parentheses refer to outer shell statistics.
<sup>b</sup> CC<sub>1/2</sub> = Pearson Correlation Coefficient between two random half datasets.
<sup>c</sup> R<sub>sym</sub> = $\sum_{hkl} \sum_i |I_{hkl,i} - \langle I_{hkl} \rangle| / \sum_{hkl} \sum_i I_{hkl,i}$ , where $I_{hkl,i}$ is the scaled intensity of the $i^{\text{th}}$ measurement of reflection $h, k, l$ , and $\langle I_{hkl} \rangle$ is the average intensity for that reflection.
<sup>d</sup> R<sub>pim</sub> = $\sum_{hkl} (1/(n-1))^{1/2} \sum_i |I_{hkl,i} - \langle I_{hkl} \rangle| / \sum_{hkl} \sum_i I_{hkl,i}$ , where $n$ is the redundancy.
<sup>e</sup> R<sub>cryst</sub> = $\sum_{hkl} |F_o - F_c| / \sum_{hkl} |F_o| \times 100$ , where $F_o$ and $F_c$ are the observed and calculated structure factors.
<sup>f</sup> R<sub>free</sub> was calculated as for R<sub>cryst</sub>, but on a test set of 5% of the data excluded from refinement.
<sup>g</sup> Calculated using MolProbity (67)

